# Crossing host boundaries: the evolutionary drivers and correlates of viral host jumps

**DOI:** 10.1101/2023.09.01.555953

**Authors:** Cedric C.S. Tan, Lucy van Dorp, Francois Balloux

**Author notes:** Correspondence: C.C.S.T. These authors contributed equally (Lucy van Dorp, Francois Balloux).

## Abstract

Most emerging and re-emerging infectious diseases stem from viruses that naturally circulate in non-human vertebrates. When these viruses cross over into humans, they can cause disease outbreaks and epidemics. While zoonotic host jumps have been extensively studied from an ecological perspective, little attention has gone into characterising the evolutionary drivers and correlates underlying these events. To address this gap, we harnessed the entirety of publicly available viral genomic data, employing a comprehensive suite of network and phylogenetic analyses to investigate the evolutionary mechanisms underpinning recent viral host jumps. Notably, we challenge conventional assumptions about the directionality of host jumps, demonstrating that humans are as much a source as a sink for viral spillover events, insofar we infer more viral host jumps from humans to other animals, than from animals to humans. Moreover, we demonstrate heightened evolution in viral lineages that involve putative host jumps. We further observe that the extent of adaptation associated to a host jump is lower for viruses with broad host ranges. Finally, we show that the genomic targets of natural selection associated to host jumps vary across different viral families with either structural or auxiliary genes being the prime targets of selection. Collectively, our results illuminate some of the evolutionary drivers underlying viral host jumps that may contribute to mitigating viral threats across species boundaries.

## Introduction

The majority of emerging and re-emerging infectious diseases in humans are caused by viruses that have jumped from wild and domestic animal populations into humans (i.e., zoonoses)^1^. Zoonotic viruses have caused countless disease outbreaks ranging from isolated cases to pandemics and have taken a major toll on human health throughout history. There is a pressing need to develop better approaches to pre-empt the emergence of viral infectious diseases and mitigate their effects. As such, there is an immense interest in understanding the correlates and mechanisms of zoonotic host jumps^1–10^.

Most studies thus far have primarily investigated the ecological and phenotypic risk factors contributing to viral host range through the use of host-virus association databases constructed mainly based on systematic literature reviews and online compendiums, including VIRION^11^ and CLOVER^12^. For example, ‘generalist’ viruses that can infect a broader range of hosts have typically been shown to be associated with greater zoonotic potential^3,5,13^. Additionally, factors such as increasing human population density^1^, alterations in human-related land use^4^, ability to replicate in the cytoplasm or being vector-borne^3^ are positively associated with zoonotic risk. However, despite global efforts to understand how viral infectious diseases emerge as a result of host jumps, our current understanding remains insufficient to effectively predict, prevent and manage imminent and future infectious disease threats. This may partly stem from the lack of integration of genomics into these ecological and phenotypic analyses.

One challenge for predicting viral disease emergence is that only a small fraction of the viral diversity circulating in wild and domestic vertebrates has been characterised to date. Due to resource and logistical constraints, surveillance studies of novel pathogens in animals often have sparse geographical and/or temporal coverage^14,15^, as they focus on selected host and pathogen taxa. Further, many of these studies do not perform downstream characterisation of the novel viruses recovered, and may lack sensitivity due to the use of PCR pre-screening to prioritise samples for sequencing^16^. As such, our knowledge of which viruses can, or are likely to emerge, and in which settings, is poor. Additionally, while genomic analyses are important for investigating the drivers of viral host jumps^17^, most studies do not incorporate genomic data into their analyses. Those that did have mostly focused on measures of host^13^ or viral^3^ diversity as predictors of zoonotic risk. As such, despite the limited characterisation of global viral diversity thus far, existing genomic databases remain a rich, largely untapped, resource to better understand the evolutionary processes surrounding viral host jumps.

Further, while research efforts have rightfully focused on zoonotic host jumps, humans are just one node in a large and complex network of host species in which viruses are endlessly exchanged, with viral zoonoses representing likely only rare outcomes of this wider ecological network. As such, viral host jumps between non-human animals remain relatively understudied. Another important process that has received less attention is human-to-animal (i.e., anthroponotic) spillover, which may impede biodiversity conservation efforts and could also negatively impact food security. For example, human-sourced metapneumovirus has caused fatal respiratory outbreaks in captive chimpanzees^18^. Anthroponotic events may also lead to the establishment of wild animal reservoirs that may re-seed infections in the human population, potentially following the acquisition of animal-specific adaptations that could increase transmissibility or pathogenicity of a virus in humans^19^. Uncovering the broader evolutionary processes surrounding host jumps across vertebrate species may therefore enhance our ability to pre-empt and mitigate the effects of infectious diseases on both human and animal health.

A major challenge for understanding macroevolutionary processes through large-scale genomic analyses is the traditional reliance on physical and biological properties of viruses to define viral taxa, which is largely a vestige of the pre-genomic era^20^. As a result, taxon names may not always accurately reflect the evolutionary relatedness of viruses, precluding robust comparative analyses involving diverse viral taxa. Notably, the International Committee on Taxonomy of Viruses (ICTV) has been strongly advocating for taxon names to also reflect the evolutionary history of viruses^20,21^. However, the increasing use of metagenomic sequencing technologies has resulted in a large influx of newly-discovered viruses that have not yet been incorporated in to the ICTV taxonomy. Furthermore, it remains challenging to formally assess genetic relatedness through multi-sequence alignments of thousands of sequences comprising diverse viral taxa, particularly for those that experience a high frequency of recombination or reassortment.

In this study, we leverage the ∼12 million viral sequences and associated host metadata hosted on NCBI to assess the current state of global viral genomic surveillance. We additionally analyse ∼59k viral sequences isolated from various vertebrate hosts, using a bespoke approach that is agnostic to viral taxonomy to understand the evolutionary processes surrounding host jumps. We ascertain overall trends in the directionality of viral host-jumps between human and non-human vertebrates and quantify the amount of detectable adaptation associated with putative host jumps. Finally, we quantify, for a subset of viruses, signatures of adaptive evolution detected in specific categories of viral proteins associated with facilitating or sustaining host jumps. Together, we provide a comprehensive assessment of potential genomic correlates underpinning host jumps in viruses across humans and other non-human vertebrates.

## Results

### An incomplete picture of global vertebrate viral diversity

Global genomic surveillance of viruses from different hosts is key to preparing for emerging and re-emerging infectious diseases in humans and animals^17,19^. To identify the scope of viral genomic data collected thus far, we downloaded the metadata of all viral sequences hosted on NCBI Virus (*n*=11,645,803; accessed 22 July 2023; **Supplementary Table 1**). Most (68%) of these sequences were associated with SARS-CoV-2, reflecting the intense sequencing efforts during the COVID-19 pandemic. Additionally, of these sequences, 93.6%, 3.3%, 1.5%, 1.1% and 0.6% were of viruses with single-stranded (ss) RNA, double-stranded (ds) DNA, dsRNA, ssDNA and unspecified genome compositions, respectively. The dominance of ssRNA viruses is not entirely explained by the high number of SARS-CoV-2 genomes, as ssRNA viruses still represent 80% of all viral genomes if SARS-CoV-2 is discounted.

Vertebrate-associated viral sequences represent 93% of this dataset, of which 93% were human-associated. The next four most sequenced vertebrates are associated to domestic animals (*Sus*, *Gallus*, *Bos*, and *Anas*) and, after excluding SARS-CoV-2, represent 15% of vertebrate viral sequences, while the other remaining genera occupy a mere 9% (**Fig. 1a**; **Extended Data Fig. 1a**), highlighting the human-centric nature of viral genomic surveillance. Further, only a limited number of non-human vertebrate families have at least ten associated viral genome sequences deposited (**Fig. 1b**), reinforcing that a substantial proportion of viral diversity in vertebrates remains uncharacterised. Viral sequences obtained from non-human vertebrates thus far also display a strong geographic bias, with most samples collected from the United States of America (USA) and China, whereas countries in Africa, Central Asia, South America and Eastern Europe are highly underrepresented (**Fig. 1c**). This geographical bias varies between the four most sequenced non-human host genera *Sus*, *Gallus*, *Anas*, and *Bos* (**Extended Data Fig. 1b**). Finally, the user-submitted host metadata associated with viral sequences, which is key for understanding global trends in the evolution and spread of viruses in wildlife, remains poor, with 45% and 37% of non-human viral sequences having no associated host information provided at the genus level, or sample collection year, respectively. The proportion of missing metadata also varies extensively between viral families and between countries (**Extended Data Fig. 2**). Overall, these results highlight the massive gaps in the genomic surveillance of viruses in wildlife globally and the need for more conscientious reporting of sample metadata.

**Figure 1.**
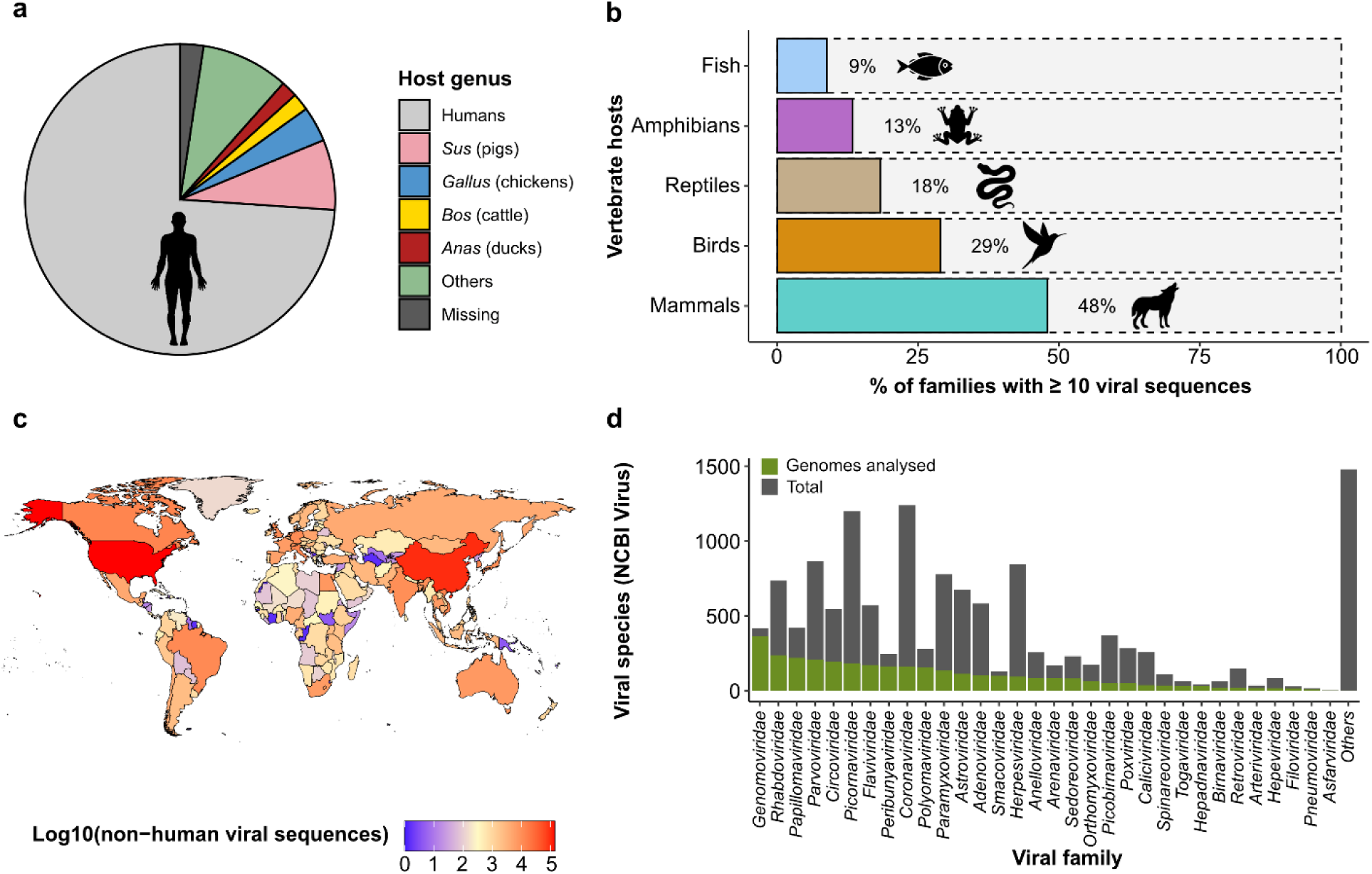
Current state of the global genomic surveillance of vertebrate viruses. (a) Proportion of non-SARS-CoV-2, vertebrate-associated viral sequences deposited on public sequence databases (*n*=2,874,732), stratified by host. Viral sequences associated with humans and the next four most-sampled vertebrate hosts are shown. Sequences with no host metadata resolved at the genus level are denoted as ‘missing’. (b) Proportion of host families represented by at least 10 associated viral sequences for the five major vertebrate host groups. (c) Global heatmap of sequencing effort, generated from all viral sequences deposited on public sequence databases that are not associated to human hosts (*n*=1,599,672). (d) Number of vertebrate viral species on NCBI Virus used for the genomic analyses in this study, stratified by viral family. The 32 vertebrate-associated viral families considered in this study are shown and the remaining 21 families that were not are denoted as ‘others’.

### Humans give more viruses to animals than they do to us

To investigate the relative frequency of anthroponotic and zoonotic host jumps, we retrieved 58,657 quality-controlled viral genomes spanning 32 viral families, associated to 62 vertebrate host orders, and representing 24% of all vertebrate viral species on NCBI Virus^22^ (**Fig. 1d**). We found that the user-submitted species identifiers of these viral genomes are poorly ascribed, with only 37% of species names consistent with those in the ICTV viral taxonomy^23^. Additionally, the genetic diversity represented by different viral species is highly variable, since they are conventionally defined based on the genetic, phenotypic and ecological attributes of viruses^20^. Thus, we implemented a species-agnostic approach based on network theory to define ‘viral cliques’ that represent discrete taxonomic units with similar degrees of genetic diversity, similar to the concept of operational taxonomic units^24^ (**Fig. 2a**; see Methods). A similar approach was previously shown to effectively partition the genomic diversity of plasmids in a biologically-relevant manner^25^. Using this approach, we identified 5,128 viral cliques across the 32 viral families that were highly concordant with ICTV-defined species (median adjusted Rand index, ARI=83%, adjusted mutual information, AMI=75%), and of which 95% were monophyletic (**Fig. 2a**). Some clique assignments aggregated multiple viral species identifiers, while others disaggregated species into multiple cliques (**Fig. 2b**; clique assignments for *Coronaviridae* illustrated in **Extended Data Fig. 3**). Despite the human-centric nature of genomic surveillance, viral cliques involving only animals represent 62% of all cliques, highlighting the extensive diversity of animal viruses in the global viral-sharing network (**Extended Data Fig. 4a**).

**Figure 2.**
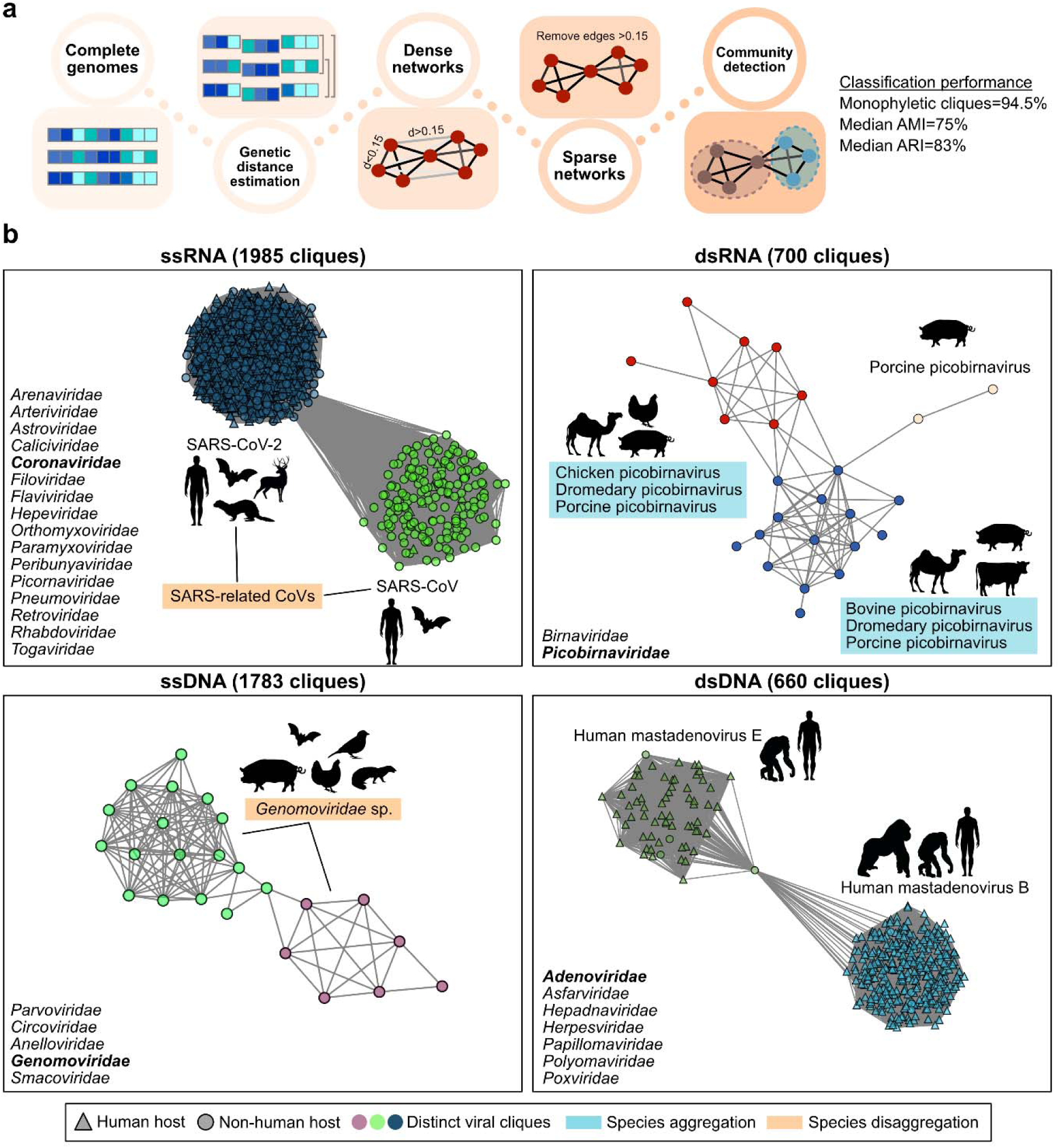
Taxonomy-agnostic approach for identifying equivalent units of viral diversity. (a) Workflow for taxonomy-agnostic clique assignments. Briefly, the alignment-free Mash^57^ distances between complete viral genomes in each viral family are computed and dense networks where nodes and edges representing viral genomes and the pairwise Mash distances, respectively, are constructed. Edges representing Mash distances >0.15 are removed to produce sparse networks, on which the community detection algorithm, Infomap^58^, is applied to identify viral cliques. Concordance with the ICTV taxonomy was assessed using the adjusted Rand Index (ARI), and adjusted mutual information (AMI). (b) Sparse networks of representative viral cliques identified within the Coronaviridae (ssRNA), Picobirnaviridae (dsRNA), Genomoviridae (ssDNA) and Adenoviridae (dsDNA). Some viral clique assignments aggregated multiple viral species, while others disaggregated species into multiple cliques. Nodes, node shapes, and edges represent individual genomes, their associated host and their pairwise Mash distances, respectively. The list of viral families considered in our analysis are shown on the bottom-left corner of each panel. Silhouettes were sourced from Flaticon.com and Adobe Stock Images (https://stock.adobe.com) with a standard license.

We then identified putative host jumps within these viral cliques by producing curated whole genome alignments to which we applied maximum-likelihood phylogenetic reconstruction. For segmented viruses, we instead used single-gene alignments as the high frequency of reassortment^26^ precludes robust phylogenetic reconstruction using whole genomes. Phylogenetic trees were rooted with suitable outgroups identified using metrics of alignment-free distances (see Methods). We subsequently reconstructed the host states of all ancestral nodes in each tree, allowing us to determine the most probable direction of a host jump for each viral sequence (approach illustrated in **Fig. 3a**). To minimise the uncertainty in the ancestral reconstructions, we considered only host jumps where the likelihood of the ancestral host state was two-fold higher than alternative host states (**Fig. 3a**; **Supplementary Methods**). Varying the stringency of this likelihood threshold yielded highly consistent results (**Extended Data Fig. 5a**), indicating that the inferred host jumps are robust to our choice of threshold. In total, we identified 12,676 viral lineages comprising 2,904 putative vertebrate host jumps across 174 of these viral cliques.

**Figure 3.**
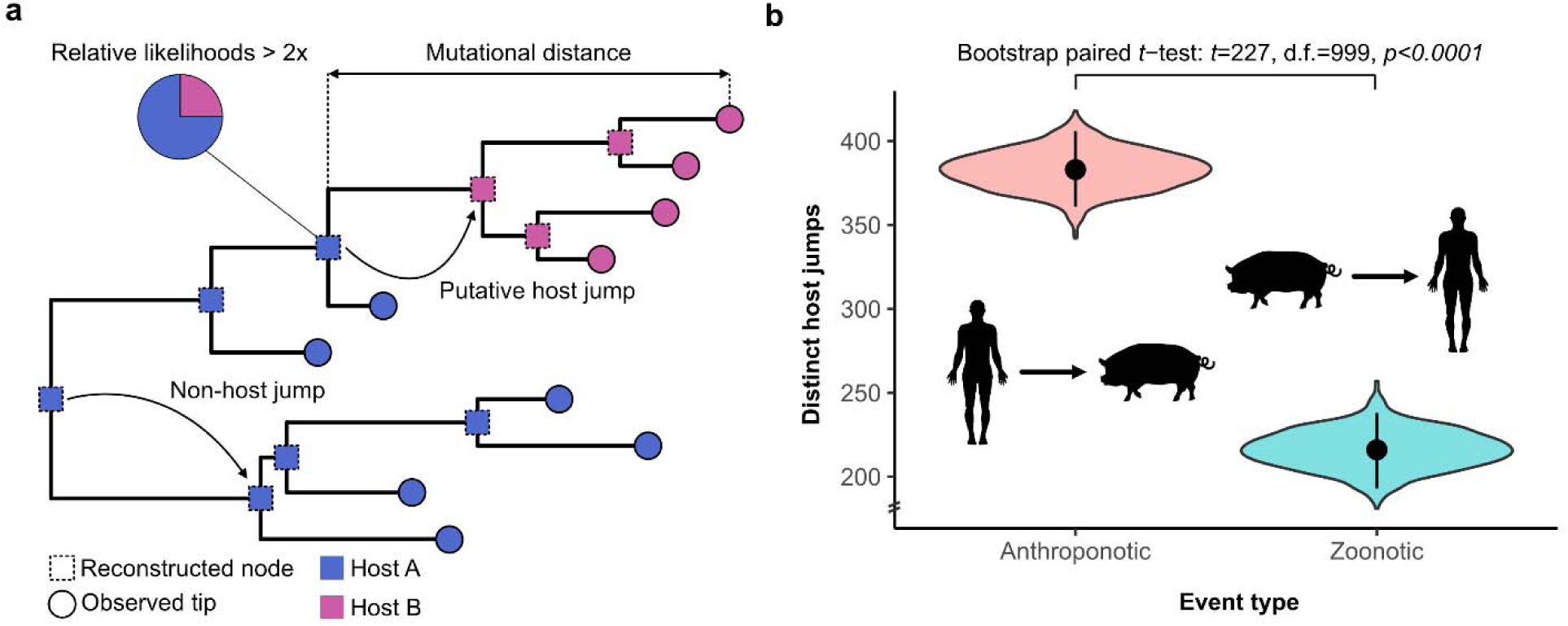
Humans give more viruses to animals than they give to us. (a) Illustration of ancestral host reconstruction approach used to identify the directionality of putative host jumps. Putative host jumps are identified if the ancestral host state has a two-fold higher likelihood than alternative host states. The mutational distance (substitutions per site) represents the sum of the branch lengths between the tip sequence and the ancestral node for which the first host state transition occurred in a tip-to-root traverse. (b) Number of distinct putative host jumps involving humans across all viral families considered (*n*=32). Black dots represent the observed point estimates for each type of host jump. The violin plots show the bootstrap distributions of these estimates, where the host jumps within each viral clique were resampled with replacement for 1000 iterations. Black lines show the 95% confidence intervals associated with these bootstrap distributions. Silhouettes were sourced from Flaticon.com and Adobe Stock Images (https://stock.adobe.com) with a standard license. A two-tailed paired t-test was performed to test for a difference in the zoonotic and anthroponotic bootstrap distributions.

Amongst the putative host jumps inferred to involve human hosts (599/2,904; 21%), we found a much higher frequency of anthroponotic compared to zoonotic host jumps (64% vs. 36%, respectively; **Fig. 3b**). This finding was statistically significant as assessed via a bootstrap paired t-test (*t*=227, d.f.=999, *p*<0.0001) and a permutation test (*p*=0.035; see Methods). Additionally, this result was robust to our choice of likelihood thresholds used during ancestral reconstruction (**Extended Data Fig. 5b**), the tree depth at which the host jump was identified (**Extended Data Fig. 5c**), and to sampling bias (**Supplementary Notes**). The highest number of anthroponotic jumps were contributed by the cliques representing SARS-CoV-2 (132/383; 34%), MERS-CoV (39/383; 10%) and Influenza A (37/383; 10%). This is concordant with the repeated independent anthroponotic spillovers into farmed, captive and wild animals, described for SARS-CoV-2^19,27–30^ and Influenza A^31,32^. Meanwhile, there has only been circumstantial evidence for human-to-camel transmission of MERS-CoV^33–35^. Noting the disproportionate number of anthroponotic jumps contributed by these viral cliques, we re-performed the analysis without them though found a significantly higher frequency of anthroponotic than zoonotic jumps (53.5% vs. 46.5%; bootstrap paired *t*-test, *t*=40, d.f.=999, *p*<0.0001), suggesting that our results are not driven solely by these cliques. Further, 16/21 of the viral families were involved in more anthroponotic than zoonotic jumps (**Extended Data Fig. 5d**), indicating that this finding is generalisable across most viruses. Overall, our results highlight the high, but largely underappreciated frequency of anthroponotic jumps amongst vertebrate viruses.

### The degree of adaptation associated to a host jump is less for multi-host viruses

Before jumping to a new host, a virus in its natural reservoir may fortuitously acquire pre-adaptive mutations that facilitate its transition to a new host. This may be followed by the further acquisition of adaptive mutations as the virus adapts to its new host environment^17^.

For each host jump inferred, we estimated the extent of both pre- and post-jump adaptations through the sum of branch lengths from the observed tip to the ancestral node where the host transition occurred (**Fig. 3a**). However, in practice, the degree of adaptation inferred may vary based on different factors, including sampling intensity and the time interval between when the host jump occurred and when the virus was isolated from its new host. As such, for each viral clique, we considered only the minimum mutational distance associated with a host jump.

We first examined whether the minimum mutational distance associated with a host jump for each viral clique was higher than the minimum for a random selection of viral lineages not involved in host jumps (**Fig. 3a**; see **Methods**). Indeed, the minimum mutational distance for a putative host jump within each clique was significantly higher than that for non-host jumps (**Fig. 4a**; one-tailed Mann-Whitney U test, *U*=6767, *p*<0.0001). Noting that both sampling intensity and the different mutation rates of viral families may confound these results, we corrected for these confounders using a logistic regression model but found a similar effect (OR_host jump_=1.31; two-tailed Z-test for slope=0, *Z*=6.58, d.f.=289, *p*<0.0001).

**Figure 4.**
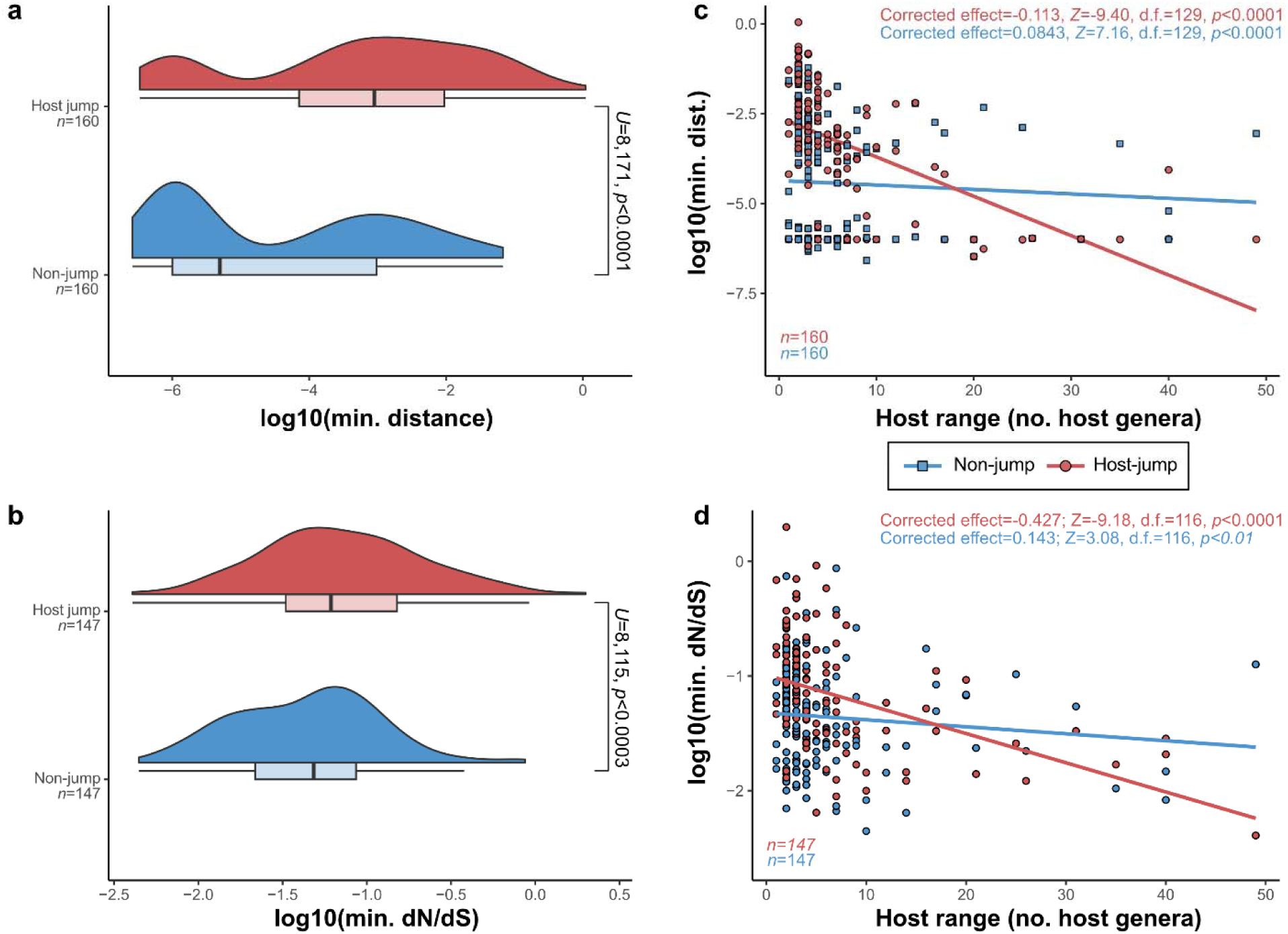
The degree of adaptation associated to host jumps is less for multi-host viruses. Distributions (Gaussian kernel densities and boxplots) of (a) minimum mutational distance and (b) minimum dN/dS for inferred host jump events and non-host jump controls on the logarithmic scale. Differences in distributions were assessed using Mann-Whitney U tests. Scatter plots of the (c) minimum mutational distance and (d) minimum dN/dS for host jump and non-host jumps. Lines represent univariate linear regression smooths fitted on the data. We corrected for the effects of sequencing effort and viral family membership using Poisson regression models. The parameter estimates in these Poisson models and their statistical significance, as assessed using two-tailed Z-tests, after performing these corrections are annotated. For all panels, each data point represents the minimum distance or minimum dN/dS across all host jump or randomly selected non-host jump lineages in a single clique. Boxplot elements are defined as follows: centre line, median; box limits, upper and lower quartiles; whiskers, 1.5x interquartile range.

We then considered the commonly used measure of directional selection acting on genomes, the ratio of non-synonymous mutations per non-synonymous site (dN) to the number of synonymous mutations per synonymous site (dS). Comparing the minimum dN/dS for host jumps within each clique, we observed that minimum dN/dS was also significantly higher for host jumps compared to non-host jumps (**Fig. 4b**; OR_host jump_=2.39; *Z*=4.84, d.f.=263, *p*<0.0001). Finally, after correcting for viral clique membership, there were no significant differences in log-transformed mutational distance (*F*(1,528)=2.23, *p*=0.136) or dN/dS estimates (*F*(1,338)=1.66, *p*=0.198) between zoonotic and anthroponotic jumps, or between forward and reverse cross-species jumps (mutational distance: *F*(1,1588)=0.538, *p*=0.463; dN/dS: F(1,1168)=0.0311, p=0.860) indicating that there are no direction-specific biases in these measures of adaptation. Overall, these results are consistent with the hypothesised need for heightened selection following a change in host environment and additionally provide confidence in our ancestral state reconstruction method for assigning host-jump status.

However, the extent of adaptive change required for a viral host-jump may vary. For instance, some zoonotic viruses may require minimal adaptation to infect new hosts while in other cases, more substantial genetic changes might be necessary for the virus to overcome barriers that prevent efficient infection or transmission in the new host. We therefore tested the hypothesis that the strength of selection associated with a host jump decreases for viruses which tend to have broader host ranges. To do so, we compared the minimum mutational distance between ancestral and observed host states to the number of host genera sampled for each viral clique. We found that the observed host range for each viral clique is positively associated with greater sequencing intensity (i.e., number of viral genomes in each clique; Pearson’s *r*=0.486; two-tailed t-test for *r*=0, *t*=34.9, d.f.=3,932, *p*<0.0001), similar to the strong correlation between per-host viral diversity and surveillance effort reported in previous studies^3,8,13^. After correcting for both sequencing effort and viral family membership, we found that the mutational threshold for host jumps tends to decrease with broader host ranges (Poisson regression, slope=-0.113; two-tailed Z-test for slope=0, *Z*=-9.40, d.f.=129, *p*<0.0001). In contrast, the relationship between mutational threshold and host range for viral lineages which have not experienced host jumps is only weakly positive (slope=0.0843; *Z*=7.16, d.f.=127, *p*<0.0001) (**Fig. 4c**). Similarly, the minimum dN/dS for a host jump decreases more substantially for viral cliques with broader host ranges (slope=-0.427; *Z*=-9.18, d.f.=116, *p*<0.0001), than for non-host jump controls (slope=0.143; *Z*=3.08, d.f.=116, *p*<0.01) (**Fig. 4d**). These trends in mutational and dN/dS thresholds were consistent when the same analysis was performed for ssDNA, dsDNA, +ssRNA and -ssRNA viruses separately (**Extended Data Fig. 6**). These results indicate that, on average, ‘generalist’ multi-host viruses experience lower degrees of adaptation when jumping into new vertebrate hosts.

### Host jump-associated adaptations are gene and region specific and vary by viral family

We next examined whether genes with different established functions displayed distinctive patterns of adaptive evolution linked to host jump events. Since gene function remains poorly characterised in the large and complex genomes of double-stranded DNA viruses, we focused on the shorter single-stranded (ss) RNA and ssDNA viral families. We selected for analysis the four non-segmented viral families with the greatest number of host jump lineages in our dataset: *Coronaviridae* (+ssRNA; *n*=2,537), *Rhabdoviridae* (-ssRNA; *n*=1,097), *Paramyxoviridae* (-ssRNA; *n*=787), and *Circoviridae* (ssDNA; *n*=695). For these viral families, we extracted all annotated protein-coding regions from their genomes and categorised them as either being associated with cell entry (termed ‘entry’), viral replication (‘replication-associated’), or virion formation (‘structural’), and classifying the remaining genes as ‘auxiliary’ genes.

For the *Coronaviridae*, *Paramyxoviridae* and *Rhabdoviridae*, the entry genes encode surface glycoproteins that could also be considered structural, but were not categorised as such given their important role in mediating cell entry. The capsid gene of circoviruses, however, encodes the sole structural protein that is also the key mediator of cell entry and was therefore categorised as structural. To estimate putative signatures of adaptation in relation to lineages which have experienced host jumps for the different gene categories, we modelled the change in log10(dN/dS) in host jumps versus non-host jumps using a linear model, while correcting for the effects of clique membership (see Methods). Contrary to our expectation that entry genes would generally be under the strongest adaptive pressures during a host jump, we found that the strength of adaptation signals for each gene category varied by family. Indeed, the strongest signals were observed for structural proteins in coronaviruses (effect=0.375, two-tailed t-test for difference in parameter estimates, *t*=4.35, d.f.=10,121, *p*<0.0001) and auxiliary proteins in paramyxoviruses (effect=0.439, *t*=2.15, d.f.=4,225, *p*=0.02) (**Fig. 5**). Meanwhile no significant adaptive signals were observed in the entry genes of all families (minimum *p*=0.3), except for the capsid gene in circoviruses (effect=0.325, *t*=2.68, d.f.=1,367, *p*=0.004) (**Fig. 5**). These findings suggest that selective pressures acting on viral genomes in relation to host jumps are likely to differ by gene functions and viral families.

**Figure 5.**
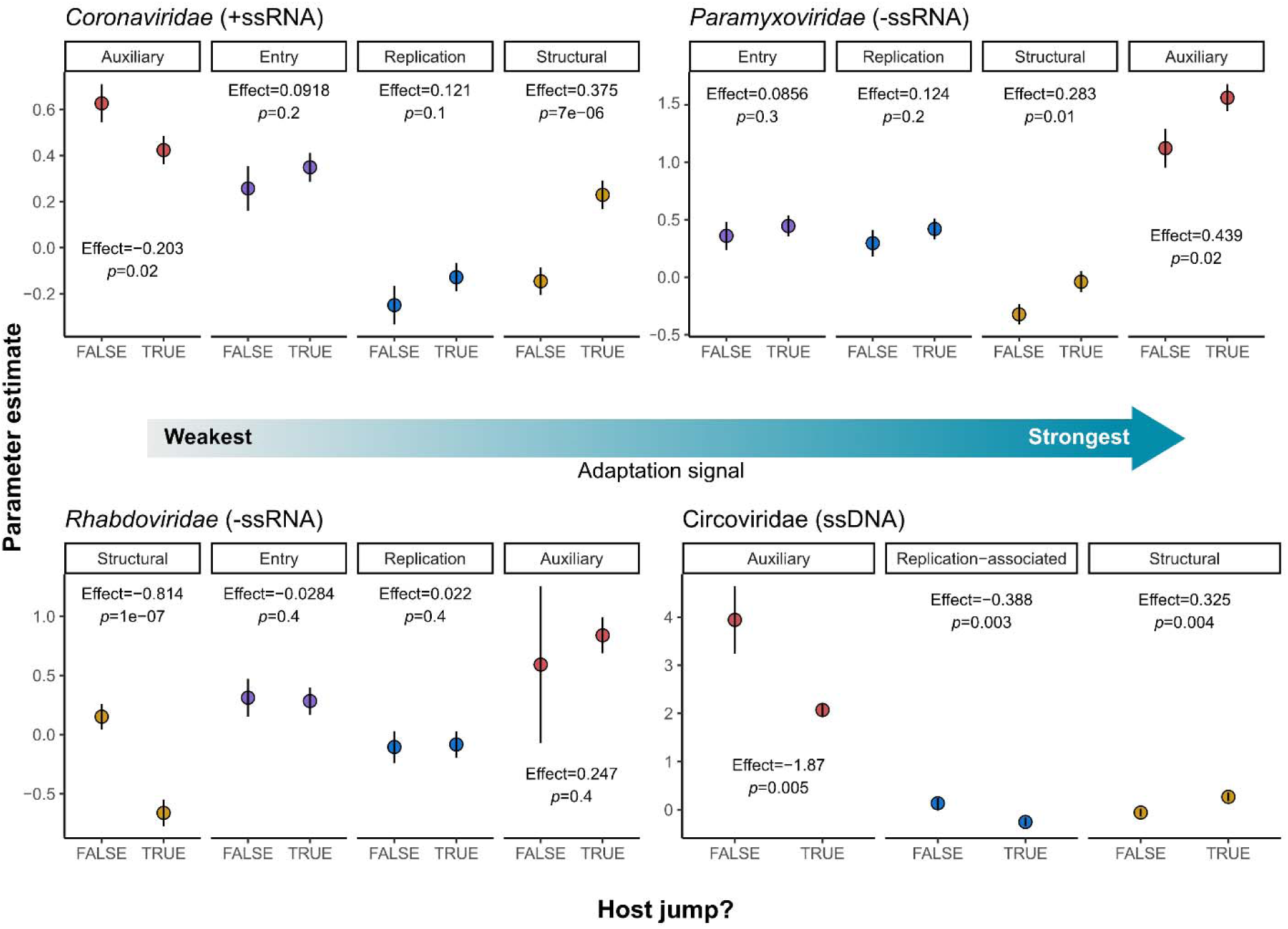
Adaptation signals are gene specific. Strength of adaptation signals per gene type were estimated using linear models. We modelled the effects of gene type and host jump status on log(dN/dS) while correcting for viral clique membership and, for each gene type, inferred the strength of adaptive signal (denoted ‘effect’) as the difference in parameter estimates for host jumps versus non-host jumps. Points and lines represent the parameter estimates and their standard errors respectively. Differences in parameter estimates were tested against zero using a one-tailed t-test. Sub-panels for each gene type were ordered from left to right with increasing effect estimates.

Given the lack of adaptive signals in the entry proteins, we further hypothesised that within each gene, adaptative changes are likely to be localised to regions of functional importance and/or that are under relatively stronger selective pressures exerted by host immunity. To test this, we focused on the spike gene (entry) of viral cliques within the *Coronaviridae* since the key region involved in viral entry is well-characterised (i.e., the receptor-binding domain (RBD))^36^. We found that dN/dS estimates consistent with adaptive evolution were indeed localised to the RBDs, but also to the N-terminal domains (NTD), of SARS-CoV-2 (genus *Betacoronavirus*), avian infectious bronchitis virus (IBV; *Gammacoronavirus*) and MERS (genus *Alphacoronavirus*) (**Extended Data Fig. 7**). This is consistent with the strong immune pressures exerted on these regions of the spike protein^37,38^ and the central role of the RBD in host-cell recognition and entry^39–41^. Overall, our results indicate that adaptive changes associated to a host jump are likely to differ by gene function, gene region and viral family.

## Discussion

The post-genomic era has opened opportunities to advance our understanding of the diversity of viruses in circulation, and the macroevolutionary principles of viral host range. Leveraging ∼59k publicly available viral sequences isolated from vertebrate hosts, we inferred that humans give more viruses to other vertebrates than they give to us across the 30 viral families we considered. We further demonstrated that host jumps are associated with heightened signals of adaptive evolution that tend to decrease in viruses with broader host ranges. This indicates that there may be a minimum mutational threshold necessary for viruses to expand their host range. Finally, we showed that adaptive evolution linked to host jumps may vary by gene function and may be localised to specific gene regions of functional importance.

To bypass the limitations of existing viral taxonomies, we used a taxonomy-agnostic approach to define roughly equivalent units of viral diversity, which formed the basis for most of the analyses presented in this study. The use of operational taxonomic units rather than traditional taxonomic species names further allowed us to perform like-for-like analyses across the entire diversity of viruses. Our approach identified cliques that were largely concordant with traditional viral species nomenclature, but also highlighted inconsistencies, where in some cases, single viral species appear to form distinct taxonomic groups while other groups of species seem to form a single group solely based on their genetic relatedness (**Fig. 2**; **Extended Data Fig. 3**). However, we do not claim that our approach should supersede existing taxonomic classification systems, especially since a robust and meaningful species definition requires the integration of viral properties with finer-scale evolutionary analyses that was not necessary for our purposes. Nevertheless, we anticipate that the use of similar network-based approaches may pave the way for the development of efficient classification frameworks that can rapidly incorporate novel, metagenomically-derived viruses into existing taxonomies.

Harnessing cliques as a mechanism of identifying clusters of related viruses for phylogenetic inspection allowed us to quantify the number and sources of recent host jump events. One important caveat to this approach is that the viral cliques involved in putative host jumps represent only a fraction of the viral diversity sequenced thus far (**Extended Data Fig. 4b**) and the patterns we observed may change as more viruses are discovered. Another caveat is that our clique assignment approach clusters viruses within approximately 15% sequence divergence, which limits our analyses to relatively recent host jump events. However, the limited divergence of the sequences within each clique also allowed us to produce more robust alignments and hence evolutionary inferences.

Of the 599 recent host jumps identified, 64% were inferred as anthroponotic (**Fig. 3b**). While the relative importance of anthroponotic versus zoonotic events has been speculated^19,32,42,43^, we provide a formal evaluation of the zoonotic-to-anthroponotic ratio in vertebrates, showing that anthroponoses are equally, if not more, critical to consider than zoonoses when assessing viral spillover dynamics. It stands to reason that the substantial global human population size and ubiquitous spatial distribution position them as a major source for viral exchange. However, it is also likely that behavioural factors might amplify the risk of anthroponotic transmission, for example through changes in land use, agricultural methods, or heightened interactions between humans and wildlife^4^. Overall, our results highlight the importance of surveying and monitoring the human-to-animal transmission of viruses.

We observed heightened evolution and adaptive signals in association with host jumps (**Fig. 4**). This result is largely intuitive, since a virus jumping into a new host is likely to be under different selective pressures exerted directly by the novel host environment and indirectly by changes in host-to-host transmission dynamics. The evolutionary signals we captured may include pre-requisite adaptations that enable a virus to infect the new host. Additionally, they likely also represent the burst of adaptive mutations which may be acquired following a host jump, which has been demonstrated for multiple viral systems^27,44–46^. Further, these signals could potentially reflect a relaxation of prior selective pressures no longer present in the novel host. We note that these signals of heightened evolution could also, in principle, be inflated by sampling bias, where two viruses circulating in the same host are more often drawn from the same population. However, this was largely controlled for in our analysis through comparisons to representative non-host jump lineages that are expected to be affected by the same sampling bias.

We observed a lower mutational and adaptive threshold associated with host jumps for viruses known to infect a broader range of hosts (**Fig. 4c** and **d**). The most likely explanation for this pattern is that some viruses are intrinsically more capable of infecting a diverse range of hosts, possibly by exploiting host-cell machinery that are conserved across different hosts. For example, sarbecoviruses (the subgenus comprising SARS-CoV-2) target the ACE2 host-cell receptor, which is conserved across vertebrates^47,48^, and the high structural conservation of the sarbecovirus spike protein^49^, may explain the observation that single mutations can enable sarbecoviruses to expand their host tropism^50^. In other words, multi-host viruses may have evolved to target more conserved host machinery that reduces the mutational barrier for them to productively infect new hosts. This may provide a mechanistic explanation for previous observations that viruses with broad host range have a higher risk of emerging as zoonotic diseases^3,5,13^.

Our approach to identifying putative host jumps hinges on ancestral state reconstruction (**Fig. 3a**), which has been shown to be affected by sampling biases^51,52^. However, we accounted for this, at least in part, by including sequencing effort as a measure of sampling bias in our statistical models, allowing us to draw inferences that were robust to disproportionate sampling of viruses in different hosts. Our approach also does not consider the epidemiology or ecology of viral transmission, as this is largely dependent on host features such as population size, social structure and behaviour for which comprehensive datasets at this scale are not currently available. We anticipate that future datasets that integrate ecology, epidemiology and genomics may allow more granular investigations of these patterns in specific host and viral systems. Additionally, the patterns we described are broad and do not capture the idiosyncrasies of individual host-pathogen associations. Those include a variety of biological features, intrinsic ones, such as the molecular adaptations required for receptor binding, as well as more complex ones including cross-immunity and interference with other viral pathogens circulating in a host population.

Overall, our work highlights the large scope of genomic data in the public domain, and its utility in exploring the evolutionary mechanisms of viral host jumps. However, the large gaps in the genomic surveillance of viruses thus far suggests that we have only just scratched the surface of the true viral diversity in nature. Additionally, despite the strong anthropocentric bias in viral surveillance, 81% of the putative host jumps identified in this study do not involve humans, emphasising the large under-appreciated scale of the global viral-sharing network (**Extended Data Fig. 8**). Widening our field of view beyond zoonoses and investigating the flow of viruses within this larger network could yield valuable insights that may help us better prepare for and manage infectious disease emergence at the human-animal interface.

## Methods

### Data acquisition, curation and quality control

The metadata of all partial and complete viral genomes was downloaded from NCBI Virus (https://www.ncbi.nlm.nih.gov/labs/virus/vssi/#/)^22^ on 22 July 2023, with filters excluding sequences isolated from environmental sources, lab hosts, or associated with vaccine strains or proviruses (*n*=11,645,803). Where possible, host taxa names in the metadata were resolved in accordance with the NCBI taxonomy^53^ using the *taxizedb* v0.3.1 package in R. User-submitted viral species names were compared to the ICTV master species list version ‘MSL38.V2’, dated 6 July 2023.

To generate a candidate list of viral sequences for further genomic analysis, the metadata was filtered to include 53 viral families known to infect vertebrate hosts based on information provided in the 2022 release of the ICTV taxonomy (https://ictv.global/taxonomy)^54^, and with reference to that provided by ViralZone (https://viralzone.expasy.org/)^55^. We then retained only sequences from viral families comprising at least 100 sequences of greater than 1000nt in length. Since the sequences of segmented viral families are rarely deposited as whole genomes and since the high frequency of reassortment^26^ precludes robust phylogenetic reconstruction, we identified sequences for single genes conserved within each of these families for further analysis (*Arenaviridae*: L segment; *Birnaviridae*: ORF1/RdRP/VP1/Segment B; *Peribunyaviridae*: L segment; *Orthomyxoviridae*: PB1; *Picobirnaviridae*: RdRP; *Sedoreoviridae*: VP1/Segment 1/RdRP; *Spinareoviridae*: Segment 1/RdRP/Lambda 3). These sequences were retrieved by applying text-based pattern matching (i.e., *grepl* in R) to query the *GenBank* sequence titles. For non-segmented genomes, we retained all non-human-associated sequences, and subsampled the human-associated sequences as follows: we selected a random subsample of 1000 SARS-CoV-2 genomes of greater than 28,000nt from distinct countries, isolation sources, and with distinct collection dates. For Influenza B, we retained only human sequences with distinct country of origins, sample types, and collection dates, and hosts of isolation. For other human-associated sequences, we retained viruses with distinct species, country, isolation source and collection date information. We then downloaded the final candidate list of viral sequences (*n*=92,973) using the *ncbi-acc-download* v0.2.8 (https://github.com/kblin/ncbi-acc-download). Further quality control of the genomes downloaded was performed using *CheckV* v1.0.1^56^, retaining sequences with more than 95% completeness (for non-segmented viruses) and less than 5% contamination (for all sequences). This resulted in a final genomic dataset comprising 58,657 observations (**Supplementary Table 2**), comprising gene sequences for segmented viruses and complete genomes for non-segmented viruses. For simplicity, we will henceforth refer to the gene sequences and complete genomes as ‘genomes’.

### Taxonomy-agnostic identification of viral cliques

To identify viral cliques, we calculated the pairwise alignment-free Mash distances of genomes within each viral family via *Mash* v1.1^57^ with a *k*-mer size of 13. This *k*-mer size ensures that the probability of observing a *k*-mer by chance given the median genome length for each clique is less than 0.01. Given a genome length, *l*, alphabet, Σ={A, T, G, C}, and the desired probability of observing a *k*-mer by chance, *q*=0.01, this was computed using the formula described previously^57^:

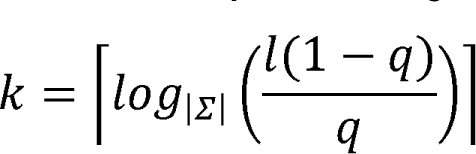

We then constructed undirected graphs for each viral family with nodes and edges representing genomes and Mash distances, respectively. We removed edges from these networks with Mash distance values greater than a certain threshold, *t*, before we applied the community-detection algorithm, Infomap^58^. This community-detection algorithm performs well in both large (>1000 nodes) and small (≤1000 nodes) undirected graphs^59^ and seeks to identify subgraphs within these undirected graphs that minimise the information required to constrain the movement of a random walker^58^. We refer to the subgraphs identified through this algorithm as ‘viral cliques’. Here, we force the community-detection algorithm to identify taxonomically-relevant cliques by removing edges with Mash distance values greater than *t*, which results in sparser graphs with closely-related genomes (e.g. from the same species) being more densely connected than more distantly-related genomes (e.g. different species). The value of *t* was selected by maximising the proportion of monophyletic cliques identified and concordance of the viral cliques identified with the viral species names from the NCBI taxonomy, based on the commonly used clustering performance metrics, adjusted mutual information (AMI) and adjusted Rand index (ARI) (**Extended Data Fig. 9**). These metrics were computed using the *AMI* and *ARI* functions in *Aricode* v1.0.2. To assess whether the viral cliques identified fulfil the species definition criterion of being monophyletic^20^, we reconstructed the phylogenies of each viral family by applying the neighbour-joining algorithm^60^ implemented in the *Ape* v5.7.1 R package on their pairwise Mash distance matrices. We then computed the proportion of monophyletic viral cliques using the *is.monophyletic* function in *Ape* v5.7.1 across the various values of *t*. Given the discordance between the NCBI and ICTV taxonomies, we applied the above optimisation protocol to *t* using the viral species names in the ICTV taxonomy. Using the NCBI viral species names, *t*=0.15 maximised both the median AMI and ARI across all families (**Extended Data Fig. 9a**), with 94.3% of the cliques identified being monophyletic (**Extended Data Fig. 9b**). Using the ICTV viral species names, *t*=0.2 and *t*=0.25 maximised the median AMI and median ARI across families (**Extended Data Fig. 9c**), with 93.7% and 87.8% cliques being monophyletic (**Extended Data Fig. 9b**), respectively. Since *t*=0.15 produced the highest proportion of monophyletic clades that were approximately concordant with existing viral taxonomies, we used this threshold to generate the final viral clique assignments for downstream analyses (**Supplementary Table 2**).

### Identification of putative host jumps

We retrieved all viral cliques that were associated with at least two distinct host genera and comprise at least 10 genomes (*n*=215). We then generated clique-level genome alignments using the *FFT-NS-2* algorithm in *MAFFT* v7.490^61,62^. We masked regions of the alignments that were poorly aligned or prone to sequencing-error by replacing alignment sites that had more than 10% of gaps or ambiguous nucleotides with N’s. Clique-level genome alignments that had more than 20% of the median genome length masked were considered to be poorly aligned and removed from further analysis (*n*=6; **Extended Data Fig. 10**). Following this procedure, we reconstructed maximum-likelihood phylogenies for each viral clique with *IQ-Tree* v2.1.4-beta^63^, using 1000 ultrafast bootstrap (UFBoot)^64^ replicates. The optimal substitution model for each tree was automatically determined using the *ModelFinder*^65^ utility native to *IQ-Tree*. To estimate the root position for each clique tree, we reconstructed neighbour-joining Mash trees for each viral clique, including ten additional genomes, whose minimum pairwise Mash distance to the genomes in each tree was 0.3-0.5, as potential outgroups. The most basal tip in these neighbour-joining Mash trees were identified and used to root the maximum-likelihood clique trees. This approach, as opposed to using maximum-likelihood phylogenetic reconstruction involving the outgroups, was used as it is difficult to reliably align clique sequences with highly divergent outgroups.

To identify putative host jumps, we performed ancestral state reconstruction on the resultant rooted maximum-likelihood phylogenies with host as a discrete trait using the *ace* function in *Ape* v5.7.1. Traversing from a tip to the root node, a putative host jump is identified if the reconstructed host state of an ancestral node is different from the observed tip state and has a two-fold greater likelihood compared to alternative states, and is different from the host state of the sampled tip. Where the tip and ancestral host states were of different taxonomic ranks, we excluded putative host jumps where the ancestral host state is nested within the tip host state, or vice versa (e.g. ‘*Homo*’ and ‘*Hominidae*’). Missing host metadata were encoded as ‘unknown’ and included in the ancestral state reconstruction analysis. Host jumps involving unknown or non-vertebrate host states were excluded from further analysis. Separately, we extracted non-host jump lineages to control for any biases in our analysis approach. To do so, we randomly selected an ancestral node – where the reconstructed host state is the same as the observed tip state and has a two-fold greater likelihood than alternative host states – for each viral genome that is not involved in any putative host-jumps. For the mutational distance and dN/dS analyses, we retained only viral cliques where non-host jump lineages could be identified. A more detailed description of the host jump inference algorithm is provided in the **Supplementary Information**.

Implementation of this algorithm yields a list of all viral lineages involving a host jump (**Supplementary Table 3a**). Since multiple lineages may involve a host transition at the same ancestral node, we calculated the number of unique host jump events as the number of distinct nodes for each unique host pair. For example, the three lineages Node1 (host A)➔Tip1 (host B), Node1 (host A)➔Tip2 (host B), and Node1 (host A)➔Tip3 (host C) would be considered as two distinct host jump events, one between hosts A and B and the other between host A and C. This counting approach was used for **Fig. 3a** and **Extended Data Fig. 5**. The lists of all 2,904 distinct host jumps are provided in **Supplementary Table 3b**.

### Calculating mutational distances and dN/dS

Mutational distance and dN/dS estimates may be lineage specific, and may depend on sampling intensity. Additionally, there is a non-linear relationship between dN/dS and branch length, that is, the estimated dN/dS decreases with increasing evolutionary distance^66^. Therefore, we opted to compare the minimum adaptive signal (i.e., minimum dN/dS) associated with a host jump for each clique. For host jump lineages, mutational distances were calculated as the sum of the branch lengths between the tip sequence and the ancestral node for which the first host state transition occurred (in substitutions per site) using the *get_pairwise_distances* function in the *Castor* v1.7.10^67^ R package, which was then multiplied by the alignment length to obtain the estimated number of substitutions (**Fig. 3a**). To calculate the dN/dS estimates, we reconstructed the ancestral sequences of ancestral nodes using the *-asr* flag in *IQ-Tree*, which is based on an empirical Bayesian algorithm (http://www.iqtree.org/doc/Command-Reference). We then extracted coding regions from the clique-level masked alignments based on the user-submitted gene annotations on NCBI GenBank (in ‘gff’ format) of each viral genome. We then computed the dN/dS estimates using Li’s^68^ method implemented in the *dnastring2kaks* function from the *MSA2dist* v1.4.0 R package (https://github.com/kullrich/MSA2dist). We calculated the minimum mutational distance and dN/dS across all host jump events in each clique for our downstream statistical analyses, which, in principle, represents the minimum evolutionary signal associated with a host jump in each viral clique. For non-host jump lineages, we similarly computed the minimum mutational distance and dN/dS across the randomly selected lineages. Estimates where dN=0 or dS=0 were removed. The list of all minimum mutational distance and minimum dN/dS estimates are provided in **Supplementary Table 3c** and **d**, respectively. The dN/dS estimates for the analysis shown in **Fig. 5** are provided in **Supplementary Table 3e**.

For the coronavirus spike gene analysis (**Extended Data Fig. 7**), spike sequences were extracted from the clique-level multiple sequence alignments, with gaps trimmed to the reference sequences (Avian infectious bronchitis virus, EU714028.1; SARS-CoV-2, MN908947.3; MERS, JX869059.2). The genomic coordinates for the functional domains of the spike proteins were derived from previous studies^36,40,69^. Estimates where dN=0 or dS=0 were removed. The dN/dS estimates are provided in **Supplementary Table 3f**.

### Statistical analyses

All statistical analyses were performed using the *stats* package native to R v4.3.1. To generate the bootstrapped distributions shown in **Fig. 3b**, we randomly resampled the host jumps within each clique with replacement (1000 iterations) and performed two-tailed paired t-tests using the *t.test* function. Mann-Whitney U tests, ANOVA, linear regressions, and Poisson and logistic regressions were implemented using *wilcox.test*, *anova*, *lm* and *glm* functions, respectively.

A permutation test was performed to assess whether the higher proportion of anthroponotic versus zoonotic jumps was statistically significant. We randomly permuted the host states in each clique for 500 iterations, while preserving the number of host-jump and non-host jump lineages (illustrated in **Extended Data Fig. 11**). The p-value was calculated as the number of iterations where the permutated anthroponotic-zoonotic ratio was greater or equal to the observed ratio.

To assess the relationship between host range and adaptative signals (**Fig. 4**), we used Poisson regressions to model the expected number of host genera observed in each viral clique, λ*_host range_*. We corrected for the number of genomes in each clique, *g*, as a measure of sampling effort, and viral family membership, *v*, by including them as fixed effects in these models. These models can be formalised for mutational distance or dN/dS, *d*, with some *p* number of viral families, and residual error, ε, as:

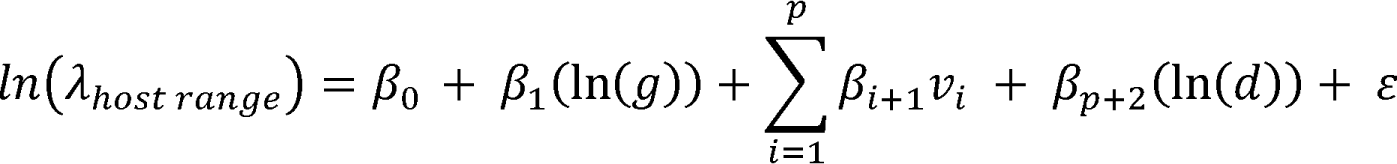

We tested if the parameter estimates were non-zero by performing two-tailed *Z*-tests implemented within the *summary* function in R.

To estimate the strength of adaptive signals for corona-, paramyxo-, rhabdo- and circoviruses (**Fig. 5**) by gene type, we implemented two linear regression models for each viral family. Since the overall adaptive signal may differ for each viral clique, we corrected for this effect using an initial linear model – where the number of viral cliques, viral clique membership, and residual, are given by *q*, *c*, and ε, respectively – as follows:

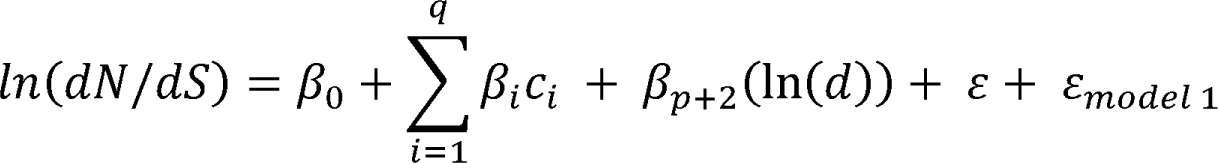

Subsequently, we used the corrected log-dN/dS estimates represented by the residuals of model 1, ε*_model 1_*, in a second linear model partitioning the effects of gene type by host jump status, *j*. Given *r* number of gene types, this model can be formalised as follows:

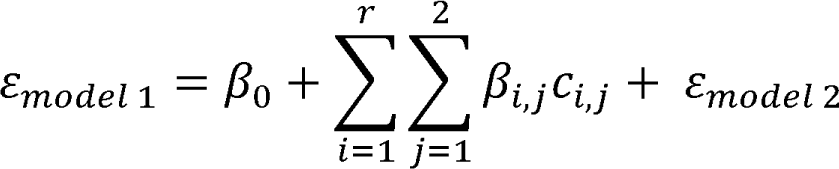

The estimated effects shown in **Fig. 5**, representative of the difference in adaptive signals associated to jump and non-host jump lineages for each gene type, was then computed as:

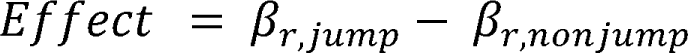

To test whether this effect is statistically significant, we used a one tailed *t*-test, with a *t* statistic computed using the standard error of the parameter estimates in model 2:

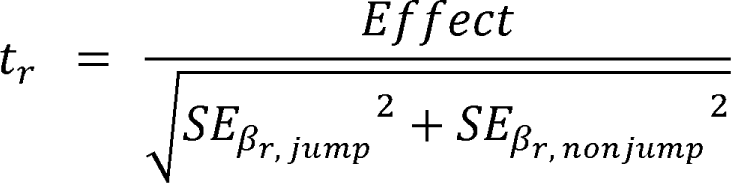

The residuals of model 2 were confirmed to be approximately normal by visual inspection (**Extended Data Fig. 12**).

### Data analysis and visualisation

All data analyses were performed using R v4.3.1. All visualisations were performed using ggplot v3.4.2^70^ or *ggtree* v3.8.2^71^. UpSet plots were created using the R package, UpSetR v1.4.0^72^.

## Acknowledgments

The authors would like to thank Rory J. Gibbs, Gemma Murray and Liam P. Shaw for their helpful feedback and discussions. C.C.S.T. is funded by the National Science Scholarship from the Agency for Science, Technology and Research (A*STAR), Singapore. F.B. and L.v.D. are funded by the European Commission (Horizon 2021-2024, END-VOC Project). L.v.D. is also funded by the UCL Excellence Fellowship. Views and opinions expressed are however those of the authors only and do not necessarily reflect those of the European Union or the European Health and Digital Executive Agency. For the purpose of open access, the corresponding author has applied a ‘Creative Commons Attribution’ (CC BY) licence to any Author Accepted Manuscript version arising. The authors acknowledge the use of the UCL Myriad High Performance Computing Facility (Myriad@UCL), and associated support services, in the completion of this work.

## Author contributions

C.C.S.T performed all analyses. L.v.D. and F.B. jointly supervised the study. C.C.S.T, L.v.D. and F.B wrote the manuscript.

## Data and code availability

All custom code used to perform the analyses reported here are hosted on GitHub (https://github.com/cednotsed/vertebrate_host_jumps). The full list of accessions considered in this study are provided in **Supplementary Table 1**. The data used for the main analyses are provided in **Supplementary Table 3a-f**. All reconstructed maximum likelihood trees and ancestral sequences used for the analyses are hosted on Zenodo (https://doi.org/10.5281/zenodo.10214868)^73^.

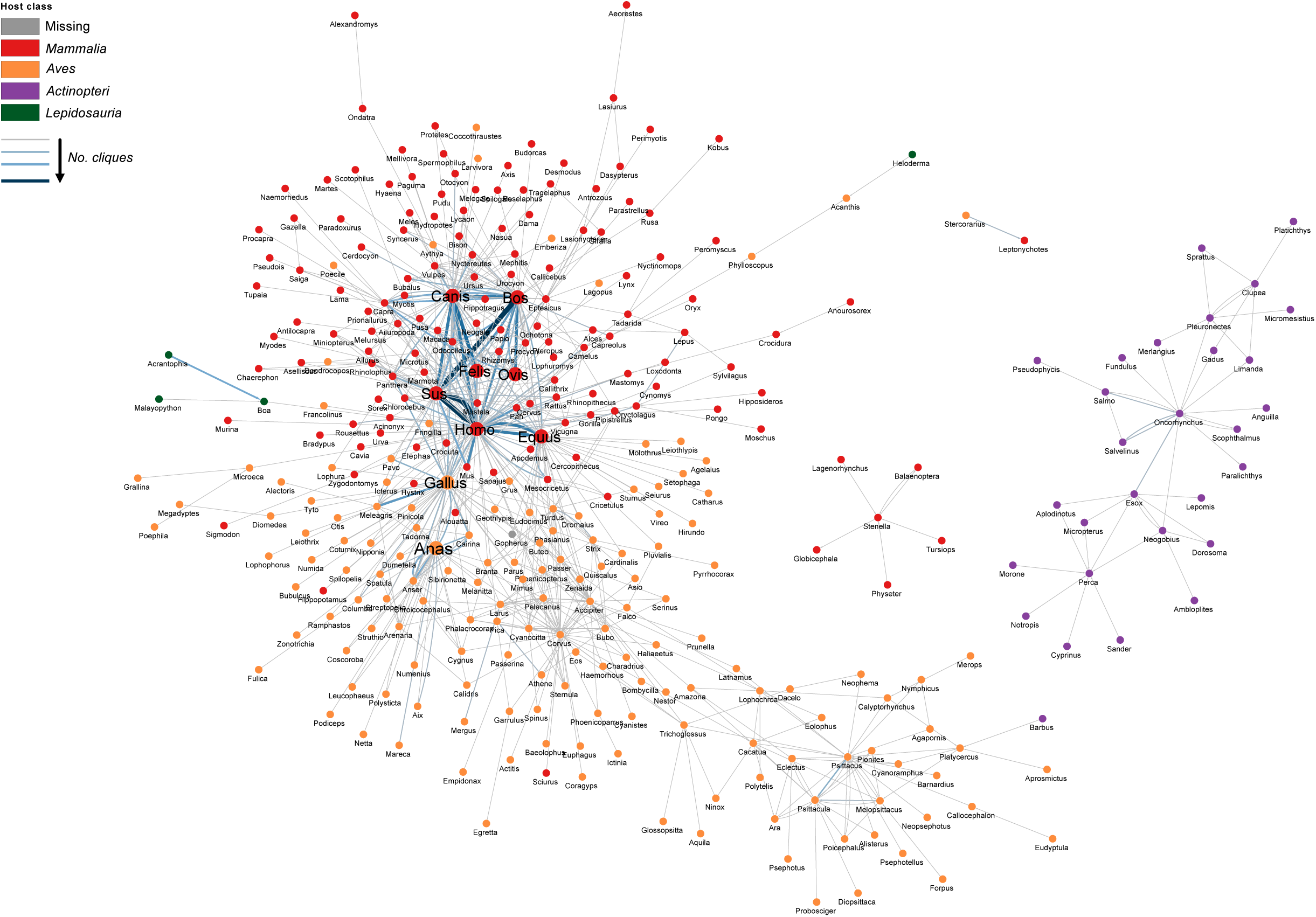

