## Supplementary Information for "Crossing host boundaries: the evolutionary drivers and correlates of viral host jumps"

* These authors contributed equally.

Supplementary Notes

*Higher frequency of anthroponotic than zoonotic jumps is robust to sampling biases*

Our estimates of the frequency of anthroponotic and zoonotic jumps across vertebrate-associated viral families rely on ancestral state reconstruction, which we note may be confounded by sampling biases. Given the high proportion of human sequences in our dataset (**Fig. 1a**), a main concern is that the likelihood of an ancestral node being reconstructed as human may be high, leading to an overestimation of the anthroponotic frequency. To assess the extent of this bias, we investigated how the estimated ratio of anthroponotic to zoonotic jumps is affected by the proportion of human-associated genomes in ‘Coronaviridae_12’, the viral clique comprising SARS-CoV-2 sequences. A high frequency of anthroponotic jumps, especially to farmed minks (*Neovison vison*)^1–3^ and wild white-tailed deer (*Odocoileus virginianus*)^4–6^ have been described. In fact, natural and experimental infections of SARS-CoV-2 have been documented for species from nearly every mammalian order. Meanwhile, only sporadic cases of animal-to-human transmission have been described, such as from pet hamsters in Hong Kong^7^, farmed mink^2^, white-tailed deer^8^, and a captive lion^9^. The extensive surveillance of SARS-CoV-2 in humans and animals therefore presents the current best case study to determine the effects of sampling biases on our results, with the expectation that humans transmit SARS-CoV-2 more frequently to animals than they do to us.

In our dataset, the proportion of human-associated sequences within Coronaviridae_12 is 63% (1000/1579). We randomly subsampled the number of human genomes, yielding proportions between 1.7% and the observed value of 63.3%, and performed ancestral reconstruction to determine the ratio of anthroponotic to zoonotic jumps. We found that as the proportion of human-associated genomes increases, the ratio of anthroponotic to zoonotic jump also increases (**Extended Data Fig. 13a**). Notably, however, the anthroponotic-zoonotic ratio is always greater than one (i.e., the expectation for SARS-CoV-2) when the proportion of human genomes is higher than 7.9% (**Extended Data Fig. 13a**), indicating that inference of whether humans are a greater source or sink for SARS-CoV-2 is robust even to extreme sampling biases. Extrapolating this finding to the full viral dataset, we re-calculated the overall anthroponotic-zoonotic ratio after removing viral cliques that comprise less than 15% animal-associated genomes, but still found a higher frequency of anthroponotic versus zoonotic jumps (**Extended Data Fig. 13b**). Further, there is good evidence in the literature for anthroponotic transmission of the viral species associated to 13/17 of these viral cliques (**Supplementary Table 4**). Overall, these results indicate that our finding that there are more anthroponotic than zoonotic jumps across vertebrate viruses is robust to sampling biases.

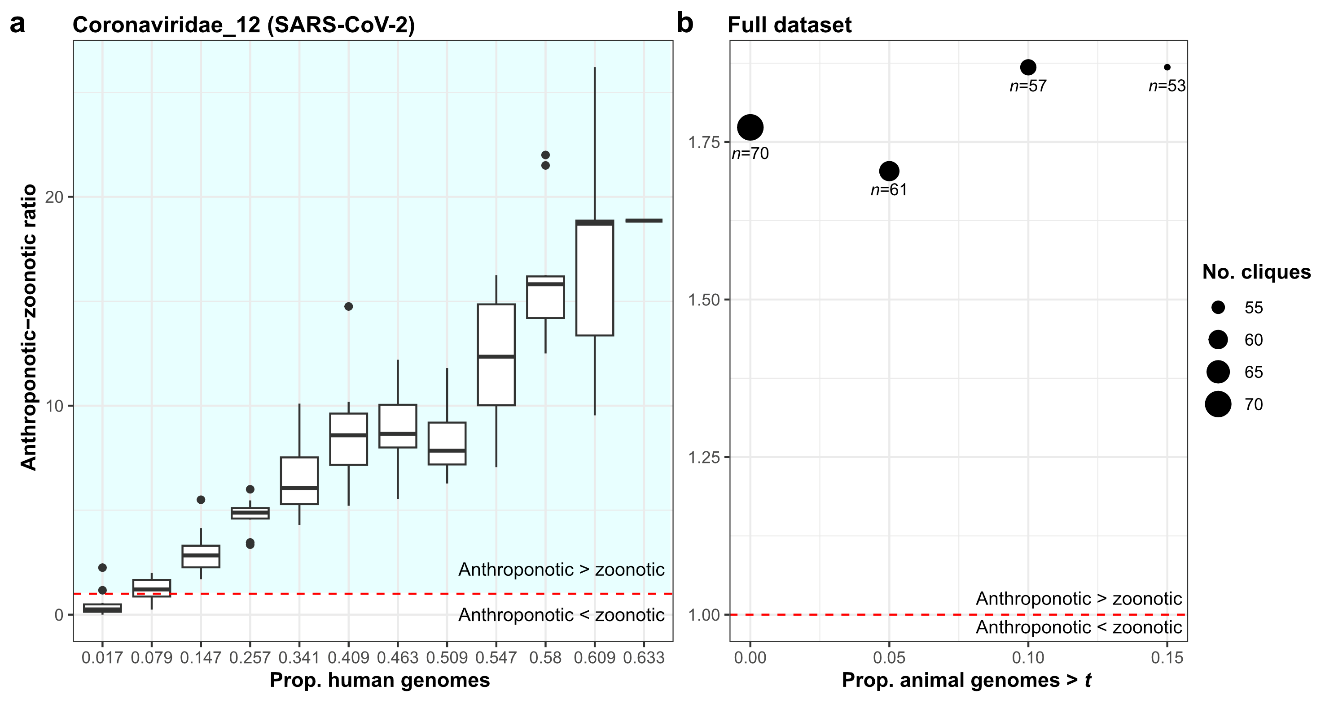

**Extended Data Figure 13**. (a) Estimates of anthroponotic-zoonotic ratio following subsampling on Coronaviridae_12, the viral clique comprising SARS-CoV-2. Human-associated genomes were randomly subsampled to various degrees prior to ancestral state reconstruction. A total of 10 iterations of the analysis per human-genome proportion were performed. Boxplot elements are defined as follows: centre line, median; box limits, upper and lower quartiles; whiskers, 1.5x interquartile range. (b) Ratio of anthroponotic to zoonotic jumps when retaining only viral cliques with greater than a certain proportion of animal-associated genomes, *t*, in the full dataset.

Supplementary Methods

*Host jump inference*

To illustrate the algorithm used for inferring putative host jumps and non-host jump lineages, we will use the clique ‘Parvoviridae_58’ that comprises mammalian bocapoviruses as an example (Extended Data Figure 14). In this clique, seven host jump lineages were identified comprising four distinct host jump events (*Sus scrofa*🡪*Rattus norvegicus*; *R. norvegicus*🡪*Mustela lutreola*; *R. norvegicus*🡪*R. rattus*; *R. flavipectus*🡪*R. norvegicus*). The algorithm used to identify host jumps traverses from tip to root, identifying the first ancestral node that differs from the host state of the tip, based on the relative likelihood of host states. For example, the algorithm traverses from the tip KY927868.1 (*R. norvegicus*) to Node 4 (*R. norvegicus*) since the likelihood of the nodes encountered so far being *R. norvegicus* is two-fold higher than alternative host states. The algorithm then traverses to Node 3 for which no host state is two-fold more likely than alternative host states, indicating that the ancestral host state for this node is ambiguous, and the traverse continues. The subsequent node is Node 2, whose host state of *S. scrofa* is two-fold more likely than alternative host states. Here, the traverse terminates, and the host jump lineage identified spans the branches between Node 2 (*S. scrofa*) and KY927868.1 (*R. norvegicus*; Supplementary Table 5).

For non-host jump lineages, the algorithm traverses from tip to node until a host state transition is encountered, which is defined as the ancestral host state being different from the tip state, and the ancestral host state having a two-fold greater likelihood than alternative states. Tips that were initially inferred to be part of a host jump lineage, or whose only ancestor is the root node, are excluded. However, by definition, any ancestral node of the same host state as the tip of interest can be chosen to represent a non-host jump lineage. For example, the branches between KY489986.1 and Node 1 or between KY489986.1 and Node 2 can be chosen, but to minimise pseudo-replication, we randomly select one ancestral node for this tip, which happens to be Node 2 in this case. The lineage leading to OM274032.1 is discarded as the next ancestral node is the root node. The full list of host jump and non-host jump lineages identified via this algorithm are shown in Supplementary Table 5.

We then calculate the number of host jumps as the number of distinct nodes where a host transition occurred for each host pair. In this case, there are seven host jump lineages comprising four distinct host jump events.

Mutational distance and dN/dS calculation

For each clique, we calculated the minimum tip-to-node distance for a host-jump and for a non-host jump lineage. The minimum mutational distance associated with a host jump and non-host jump in this case is 0.00605 and 0.000824 substitutions per site, respectively (**Supplementary Table 5**). For dN/dS, we removed all lineages where the genome-wide dN or dS value is zero. We then calculated the minimum dN/dS as a measure of the minimum extent of adaptation observed for this clique, which in this case is 0.0359 for host jump lineages. All non-host jump lineages identified here had either zero dN or dS values and so no minimum dN/dS value could be calculated for this clique. As a result, this clique was omitted from our dN/dS analyses as no non-host jump can be used as a suitable control.

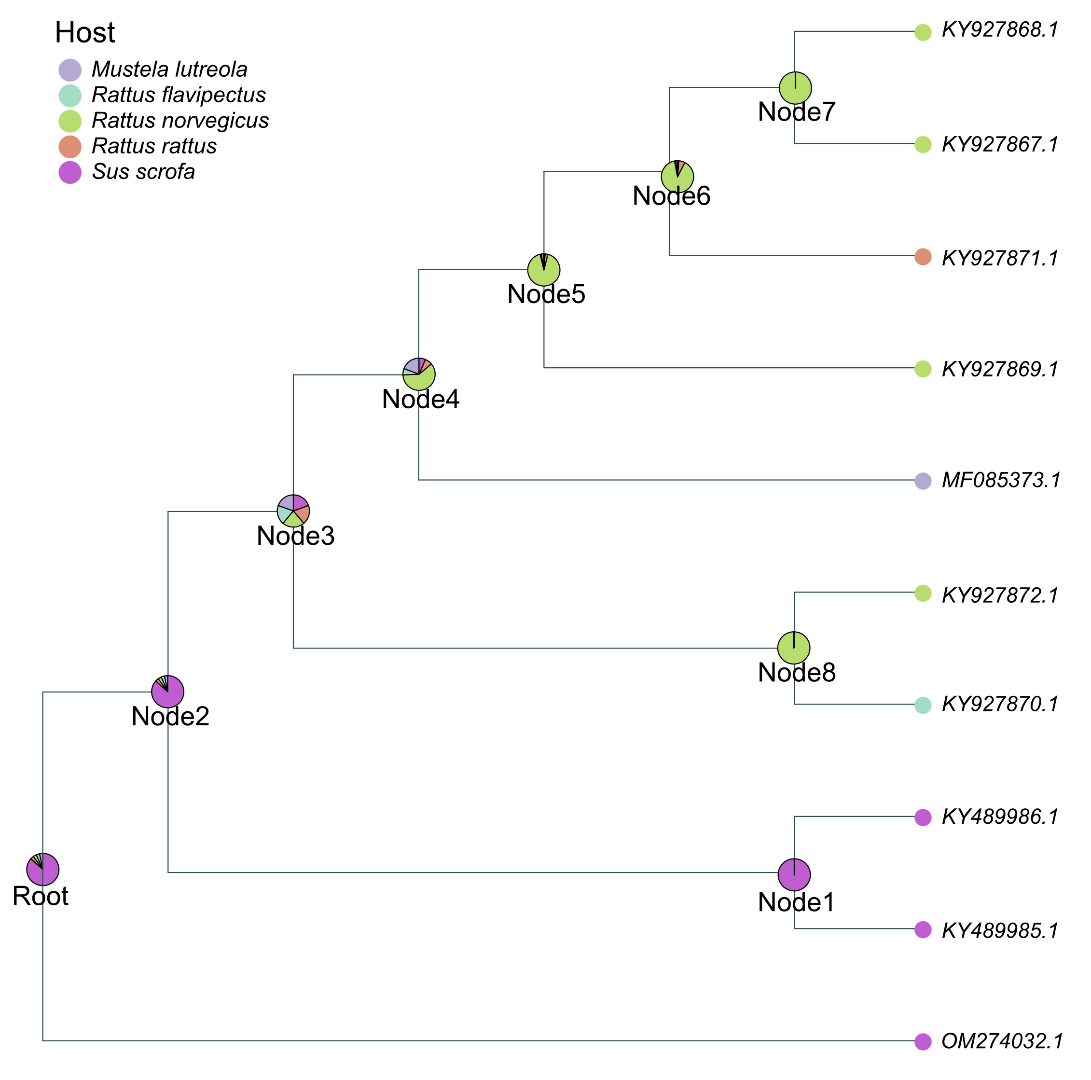

**Extended Data Figure 14**. Maximum likelihood tree for the viral clique Parvoviridae_58, rooted at OM274032.1. Piecharts provide the relative likelihoods of all host states represented in this tree, as determined by maximum-likelihood ancestral state reconstruction.

**Supplementary Table 4**: Evidence from the literature reporting natural infection of animal hosts with viral species associated to cliques with less than 15% animal-associated genomes.

| **Viral clique** | **Prop. animal genomes** | **Viral species** | **Recipient** | **Study** |
| --- | --- | --- | --- | --- |
| Orthomyxoviridae_4 | 0.002 | Influenza B virus | *Sus scrofa* | *Ran et al.*^10^ |
|  |  |  | *Rhizomys pruinosus* | *He et al.*^11^ |
| Pneumoviridae_6 | 0.01 | Orthopneumovirus hominis | *Pangolins* | Ye et al.^12^ |
| Pneumoviridae_4 | 0.01 | Orthopneumovirus hominis | *Pangolins* | Ye et al.^12^ |
| Orthomyxoviridae_11 | 0.02 | Gammainfluenzavirus influenzae | *Sus scrofa* | Kimura et al.^13^ |
| Adenoviridae_8 | 0.02 | Human mastadenovirus A | *Pan troglodytes* | Zhou et al.^14^ |
| Picornaviridae_25 | 0.02 | Enterovirus A | *Mus musculus* | Experimental infection likely |
| Picornaviridae_24 | 0.02 | Enterovirus B | *Rhinopithecus roxellana* | Tan et al.^15^ |
| Paramyxoviridae_4 | 0.03 | Respirovirus pneumoniae | *Pan troglodytes*  *Papio ursinus*  *Cercopithecus mitis* | Negrey et al.^16^ |
| Pneumoviridae_2 | 0.03 | Metapneumovirus hominis | *Gorrila beringei* | Mazet et al.^17^ |
|  |  |  | *Pan troglodytes* | Negrey et al.^16^ |
| Adenoviridae_5 | 0.06 | Human mastadenovirus C | *Gorilla gorilla* | Medkour et al.^18^ |
| Caliciviridae_36 | 0.07 | Norovirus | *Macaca mulatta* | Jiang et al.^19^ |
| Sedoreoviridae_2 | 0.08 | Rotavirus A | *Bos taurus* | Bwogi et al.^20^ |
|  |  |  | *Sus scrofa* | Bwogi et al.^20^ |
| Picornaviridae_30 | 0.09 | Salivirus A | *Pan* | Negrey et al.^21^ |
| Sedoreoviridae_1 | 0.12 | Rotavirus A | *Bos taurus* | Bwogi et al.^20^ |
|  |  |  | *Sus scrofa* | Bwogi et al.^20^ |
| Togaviridae_1 | 0.13 | Chikungunya virus | *Macaca fascicularis* | Sam et al.^22^ |
| Herpesviridae_17 | 0.14 | Human alphaherpesvirus | *Chlorocebus sabaeus* | No evidence |
| Arenaviridae_31 | 0.14 | Mammarenavirus lassaense | *Mastomys natalensis* | No evidence |
|  |  |  | *Lophuromys sikapusi* | No evidence |

**Supplementary Table 5**: Host jump and non-host jump lineages identified for the viral clique Parvoviridae_58. Minimum mutational distance and dN/dS values for this clique are underlined.

| **Ancestral node** | **Tip** | **Ancestral state** | **Observed state** | **Is host jump?** | **Mutational distance (subst./site)** | **dN** | **dS** | **dN/dS** |
| --- | --- | --- | --- | --- | --- | --- | --- | --- |
| Node2 | KY927867.1 | *S. scrofa* | *R. norvegicus* | Yes | 0.204 | 0.0433 | 0.497 | 0.0872 |
| Node2 | KY927868.1 | *S. scrofa* | *R. norvegicus* | Yes | 0.204 | 0.0433 | 0.497 | 0.0872 |
| Node2 | KY927869.1 | *S. scrofa* | *R. norvegicus* | Yes | 0.200 | 0.0428 | 0.494 | 0.0867 |
| Node2 | KY927872.1 | *S. scrofa* | *R. norvegicus* | Yes | 0.193 | 0.0435 | 0.470 | 0.0927 |
| Node6 | KY927871.1 | *R. norvegicus* | *R. rattus* | Yes | 0.00605 | 0.000510 | 0.0142 | 0.0359 |
| Node4 | MF085373.1 | *R. norvegicus* | *M. lutreola* | Yes | 0.00938 | 0.00292 | 0.0276 | 0.106 |
| Node8 | KY927870.1 | *R. norvegicus* | *R. flavipectus* | Yes | 0.0126 | 0.00409 | 0.0444 | 0.0921 |
| Node1 | KY489985.1 | *S. scrofa* | *S. scrofa* | No | 0.000824 | 0 | 0.00206 | N.A. |
| Node2 | KY489986.1 | *S. scrofa* | *S. scrofa* | No | 0.00437 | 0 | 0 | N.A. |
