## Extended Data Fig. for "Crossing host boundaries: the evolutionary drivers and correlates of viral host jumps"

* These authors contributed equally.

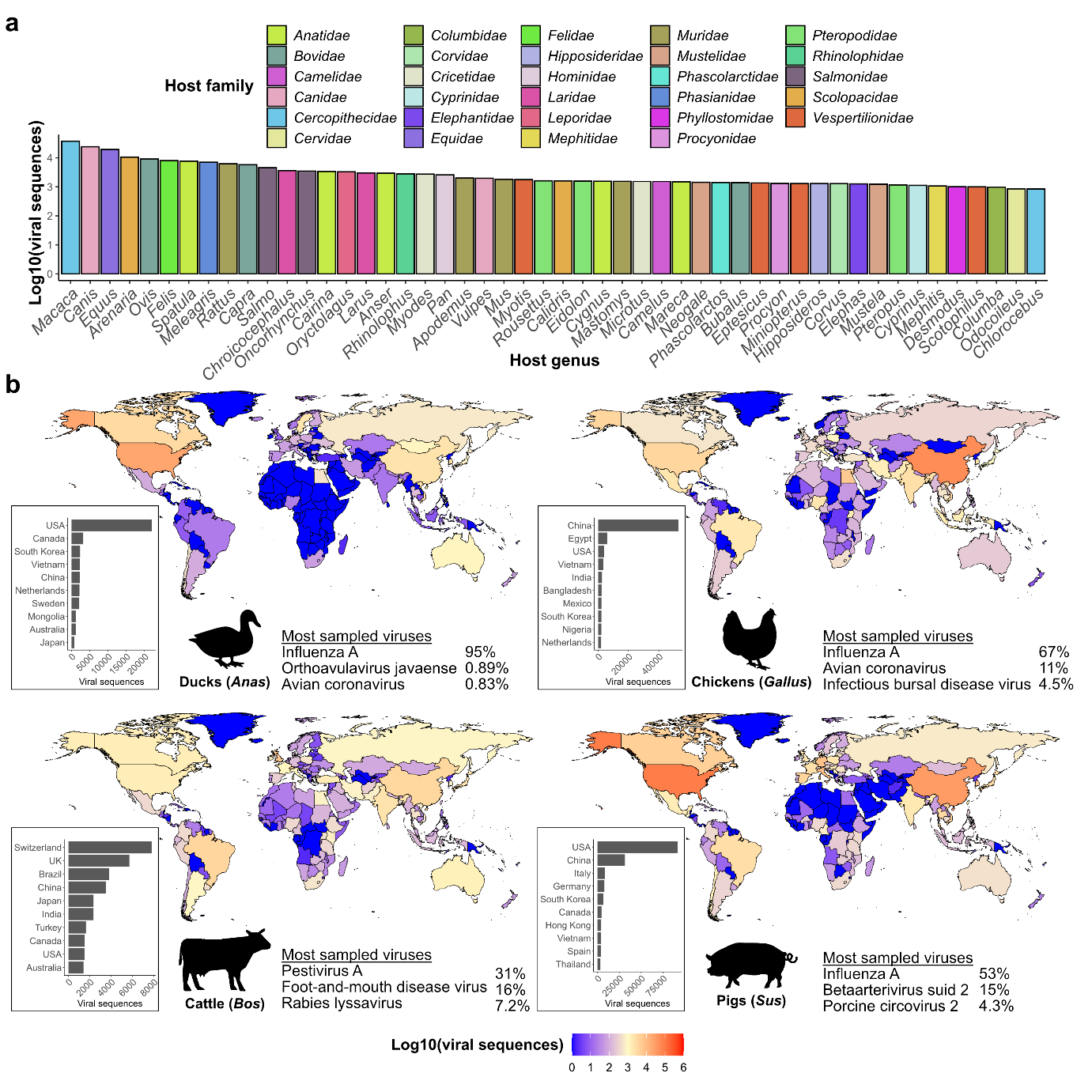

**Extended Data Figure 1**. (a) Number of viral sequences, excluding SARS-CoV-2, associated with the top 50 vertebrate hosts observed in the ‘others’ category as shown in main text Fig. 1a. (b) Number of viral sequences stratified by the four most-sequenced non-human animals, excluding SARS-CoV-2. The number of viral sequences for the top 10 countries are shown as bar plots. The percentage of viral sequences for the top three most sequenced viral species for each host are annotated.

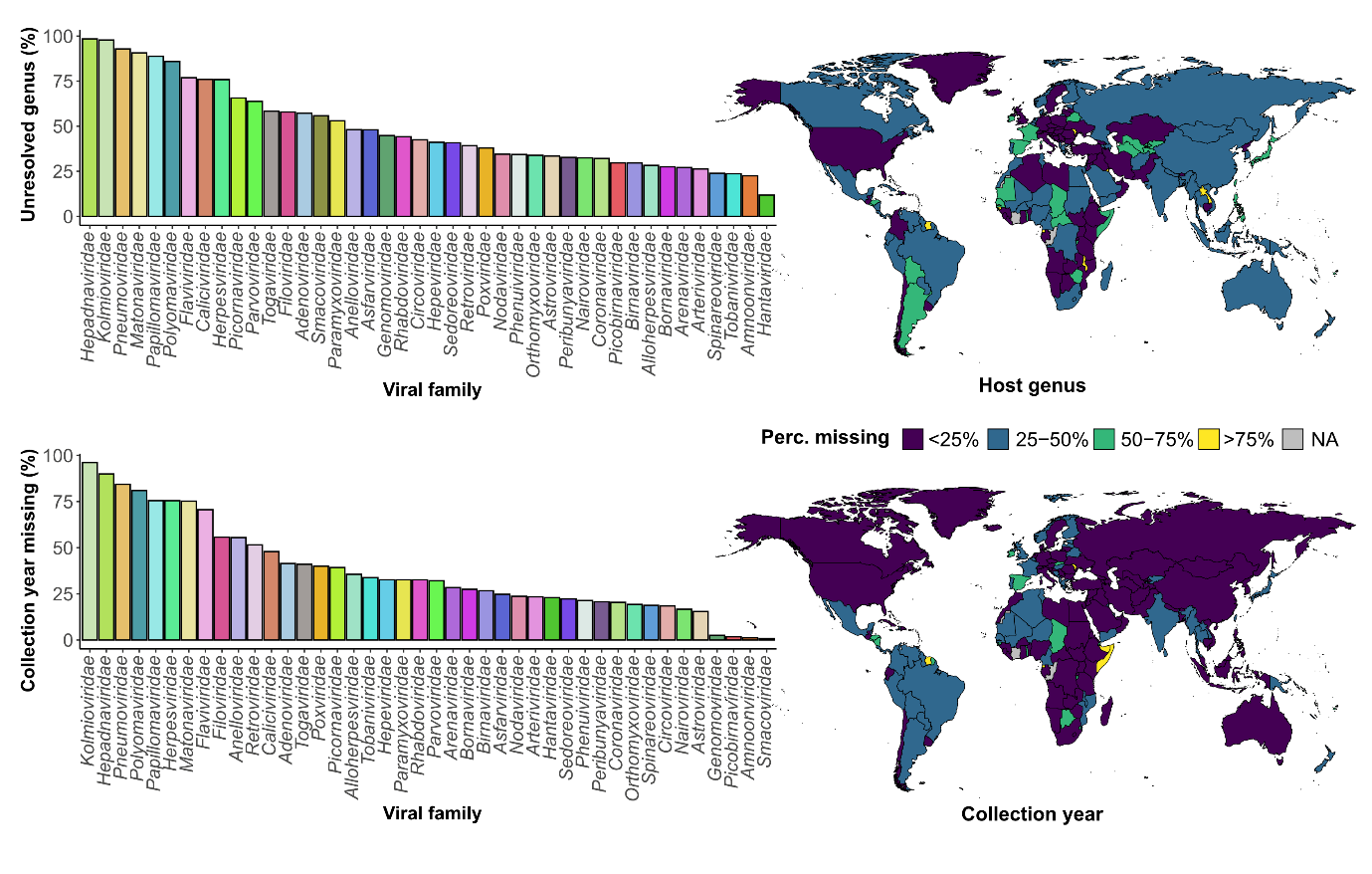

**Extended Data Figure 2**. (Top) Proportion of all viral sequences associated to non-human vertebrates (*n*=1,599,672) with missing genus information or (bottom) sample collection year, stratified by viral family or country of origin. Countries with no associated sequences are denoted ‘NA’.

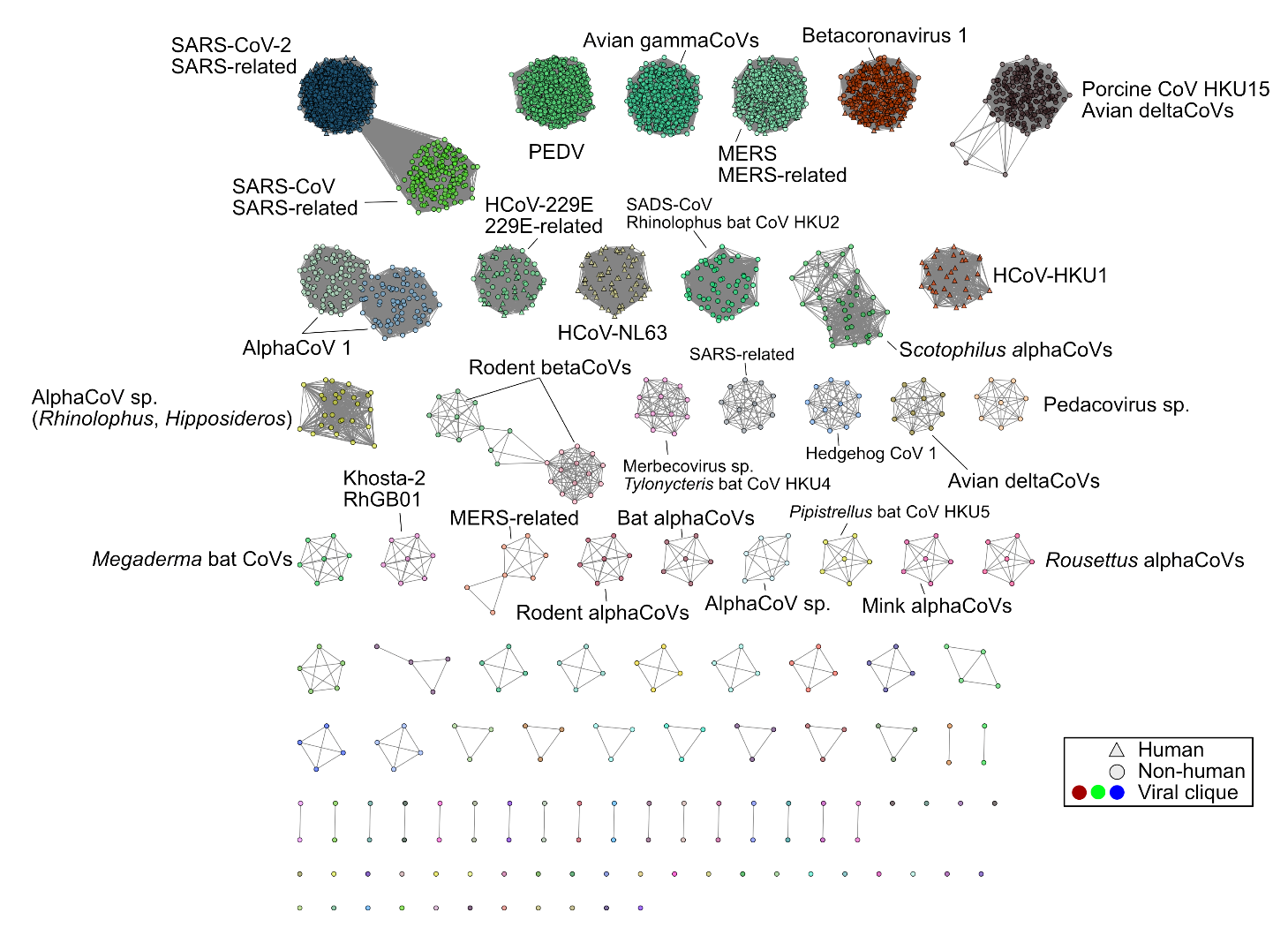

**Extended Data Figure 3**. Sparse networks of viral cliques identified (see Methods) and their corresponding user-submitted species names for the Coronaviridae, similar to main text Fig. 2. Nodes, node shapes, and edges represent individual genomes, their associated host and their pairwise Mash (alignment-free) distances, respectively.

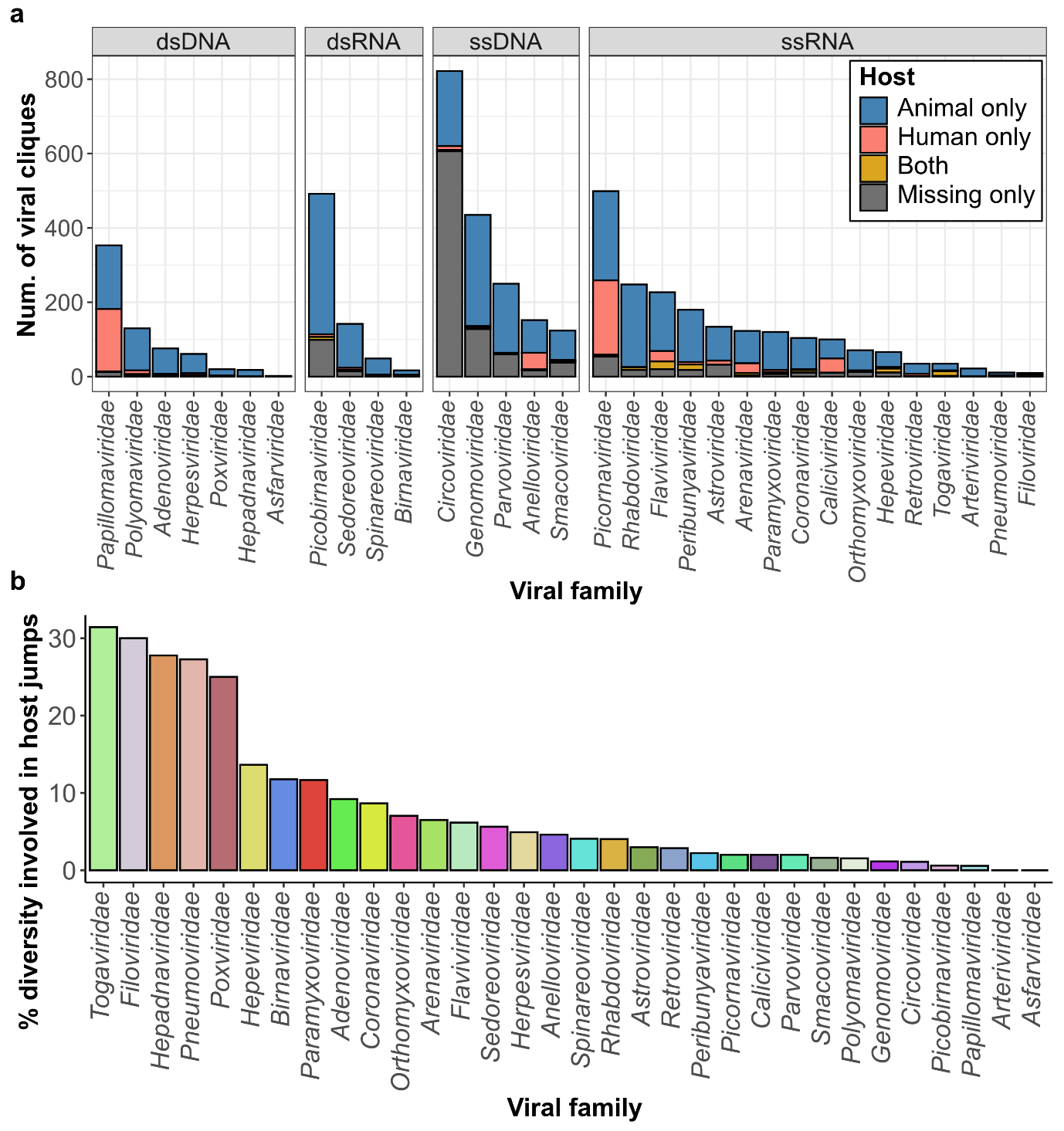

**Extended Data Figure 4**. (a) Number of viral cliques identified stratified by viral family. Cliques with only animal-associated sequences, human-associated sequences, or both are annotated. (b) Percentage of viral cliques involving at least one of the 2,904 putative host jumps inferred, stratified by viral family.

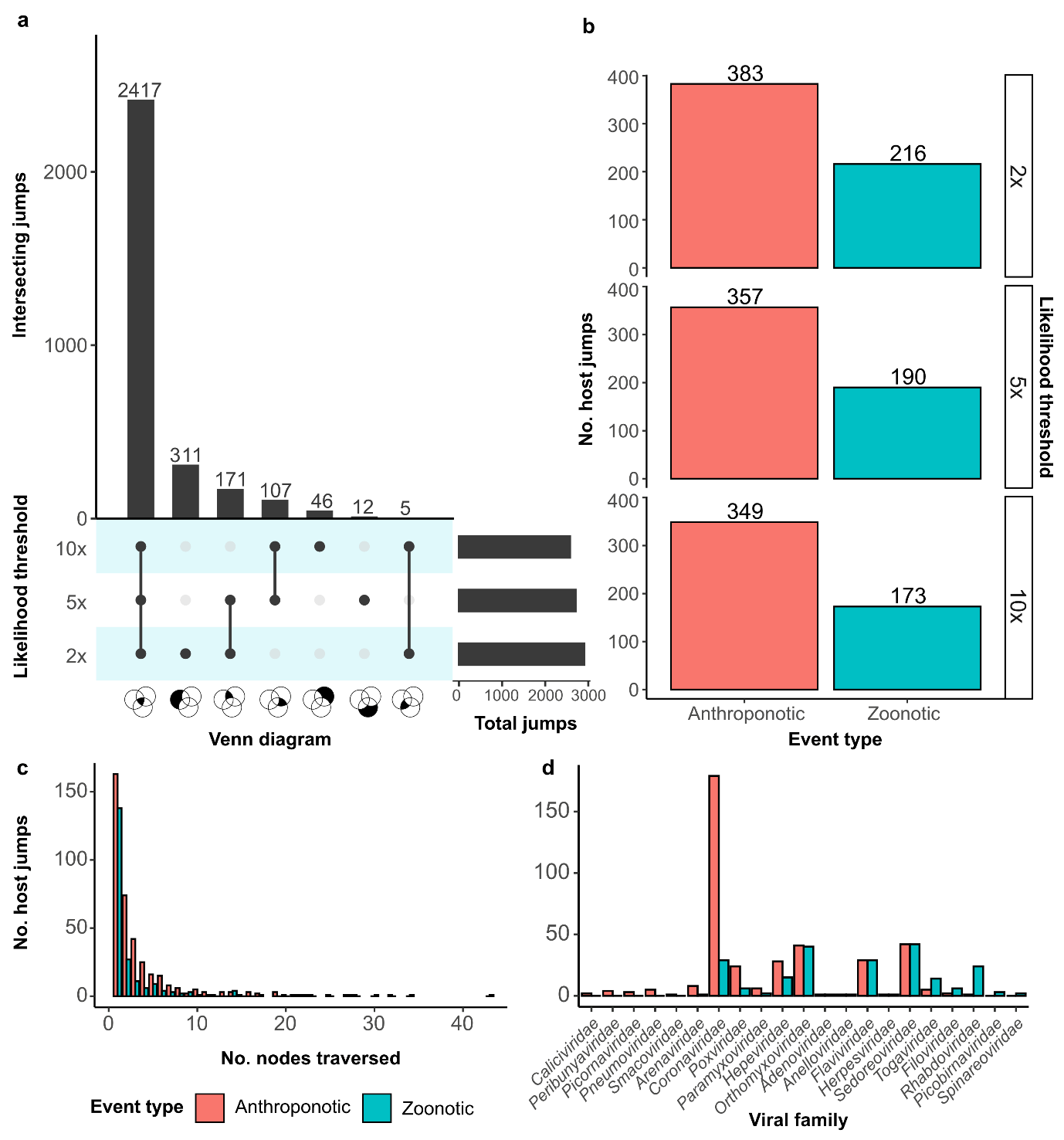

**Extended Data Figure 5**. (a) UpSet plot providing the intersecting host jumps identified via ancestral reconstruction when using a two-fold, five-fold or ten-fold likelihood threshold. (b) Bar plot showing the number of anthroponotic and zoonotic events inferred using various likelihood thresholds, (c) at different ancestral node depths, and (d) stratified by viral family. For (b), the number of anthroponotic and zoonotic host jumps were stratified by the depth of the ancestral node in the tip-to-node traversal. Since multiple host jump lineages can involve the same ancestral node, the tip-to-node depths may vary depending on which lineage is selected. As such, we randomly selected a viral lineage for each distinct host jump event for this analysis.

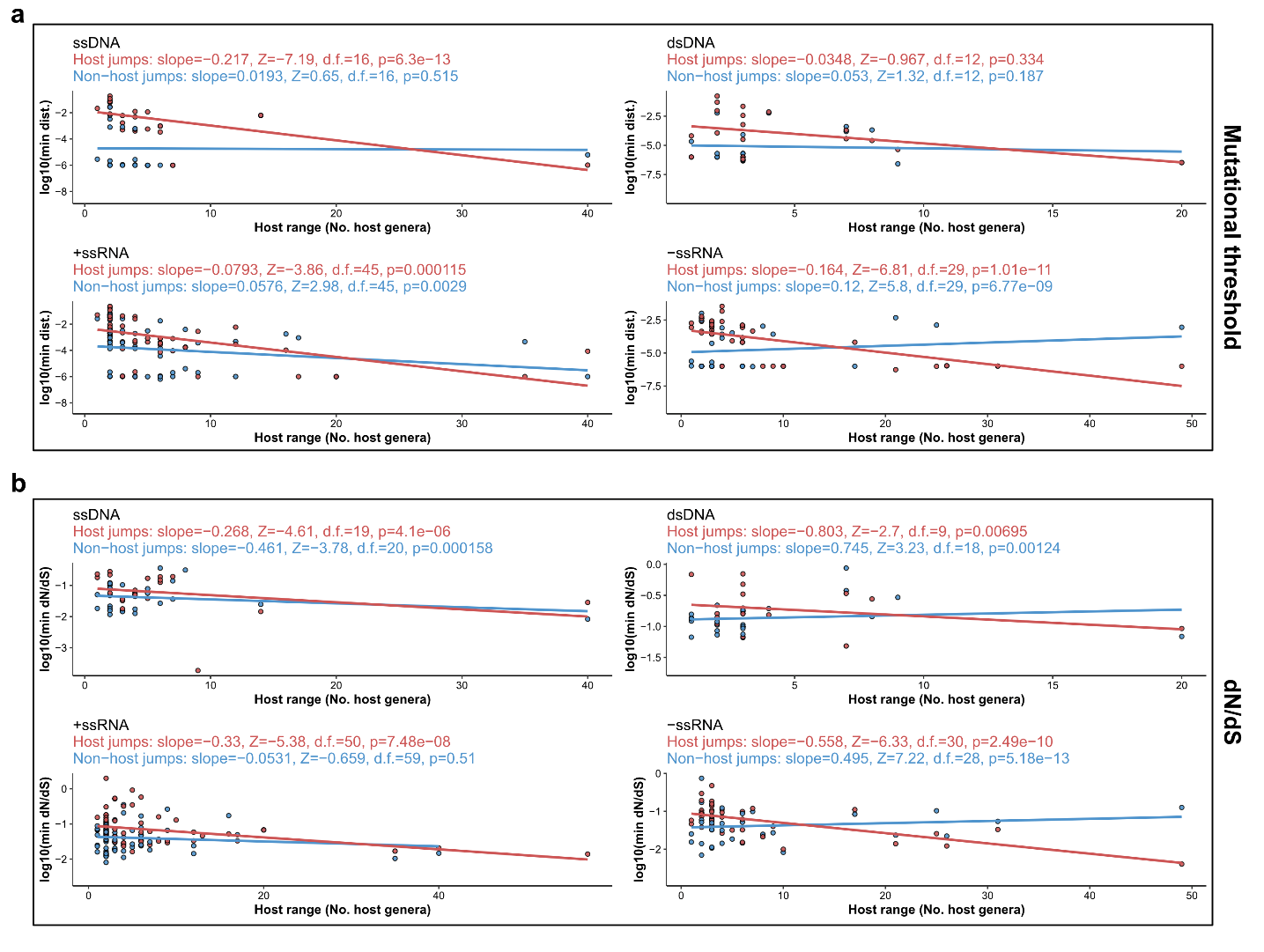

**Extended Data Figure 6**. Analysis of relationships between host range and estimated adaptive thresholds, similar to Fig. 3, but only considering ssDNA, dsDNA, +ssRNA or -ssRNA viruses. Distributions of minimum (a) mutational distance and (b) dN/dS for host jump and non-host jumps on the logarithmic scale. We corrected for the effects of sequencing effort and viral family membership using Poisson regression models. The estimated effects of patristic distance on host range after these corrections are annotated. We tested whether the estimated effects were non-zero using two-tailed Z-test. For all panels, each data point represents the minimum distance or dN/dS across all host jump or randomly selected non-host jump lineages in a single clique. Line segments represent linear regression smooths without correction.

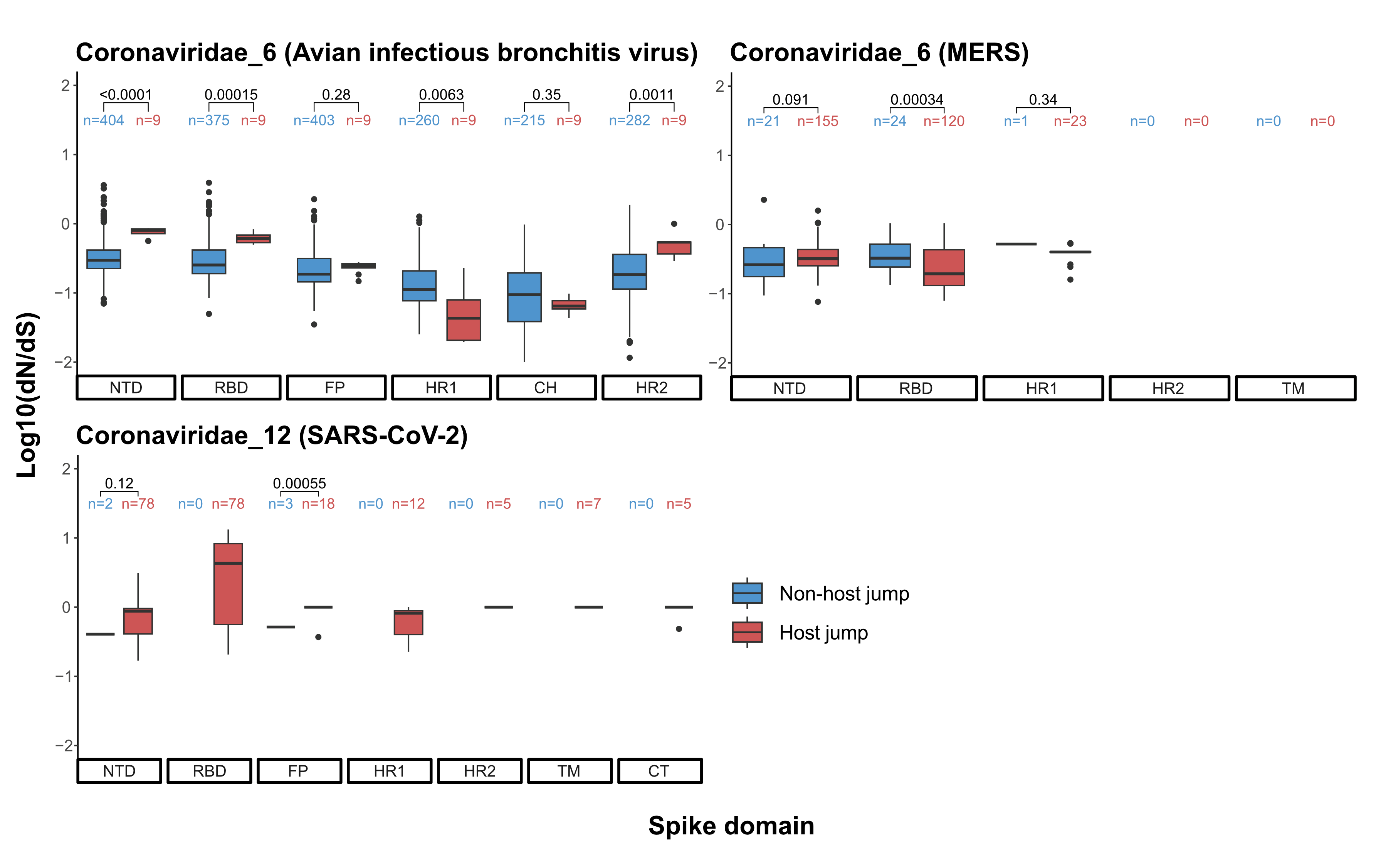

**Extended Data Figure 7**. (a) Analysis of the log10(dN/dS) estimates associated to different functional domains encoded by the coronavirus spike gene: N-terminal domain (NTD), receptor-binding domain (RBD), fusion peptide (FP), heptad repeats 1 and 2 (HR1 and HR2), central helix (CH), transmembrane (TM), C-terminal domains (CT). Estimates with dN=0 or dS=0 were removed and the remaining number of sequences for each domain and viral clique are annotated. (b) Differences in distributions were tested for using Mann-Whitney U tests and the corresponding p-values are annotated. Boxplot elements are defined as follows: centre line, median; box limits, upper and lower quartiles; whiskers, 1.5x interquartile range.

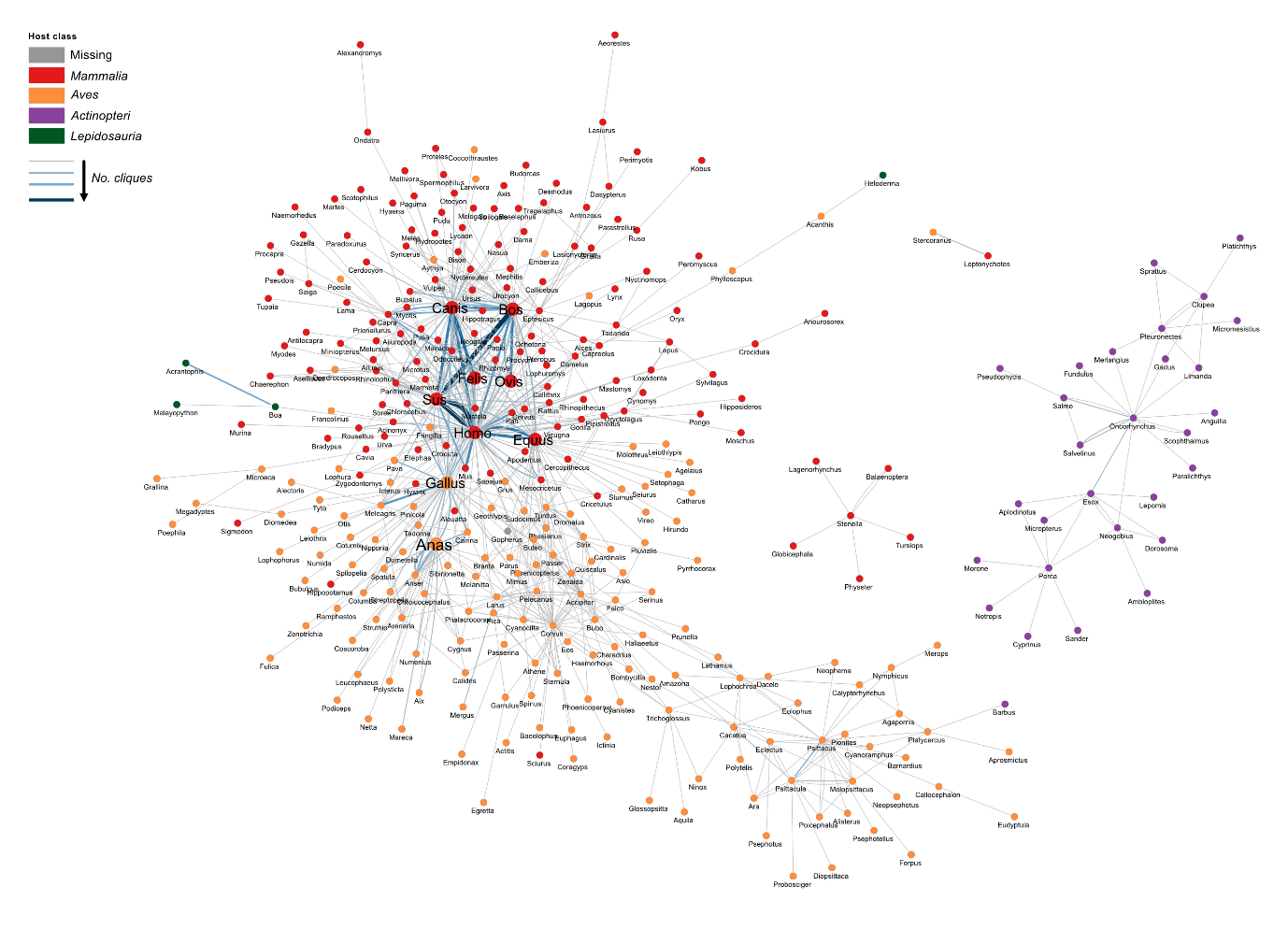

**Extended Data Figure 8**. (Also provided as separate PDF). Directed network of the vertebrate viral-sharing network, where nodes and edges represent host genera and the number of viral cliques shared. Edge widths and colour are indicative of the number of viral cliques shared.

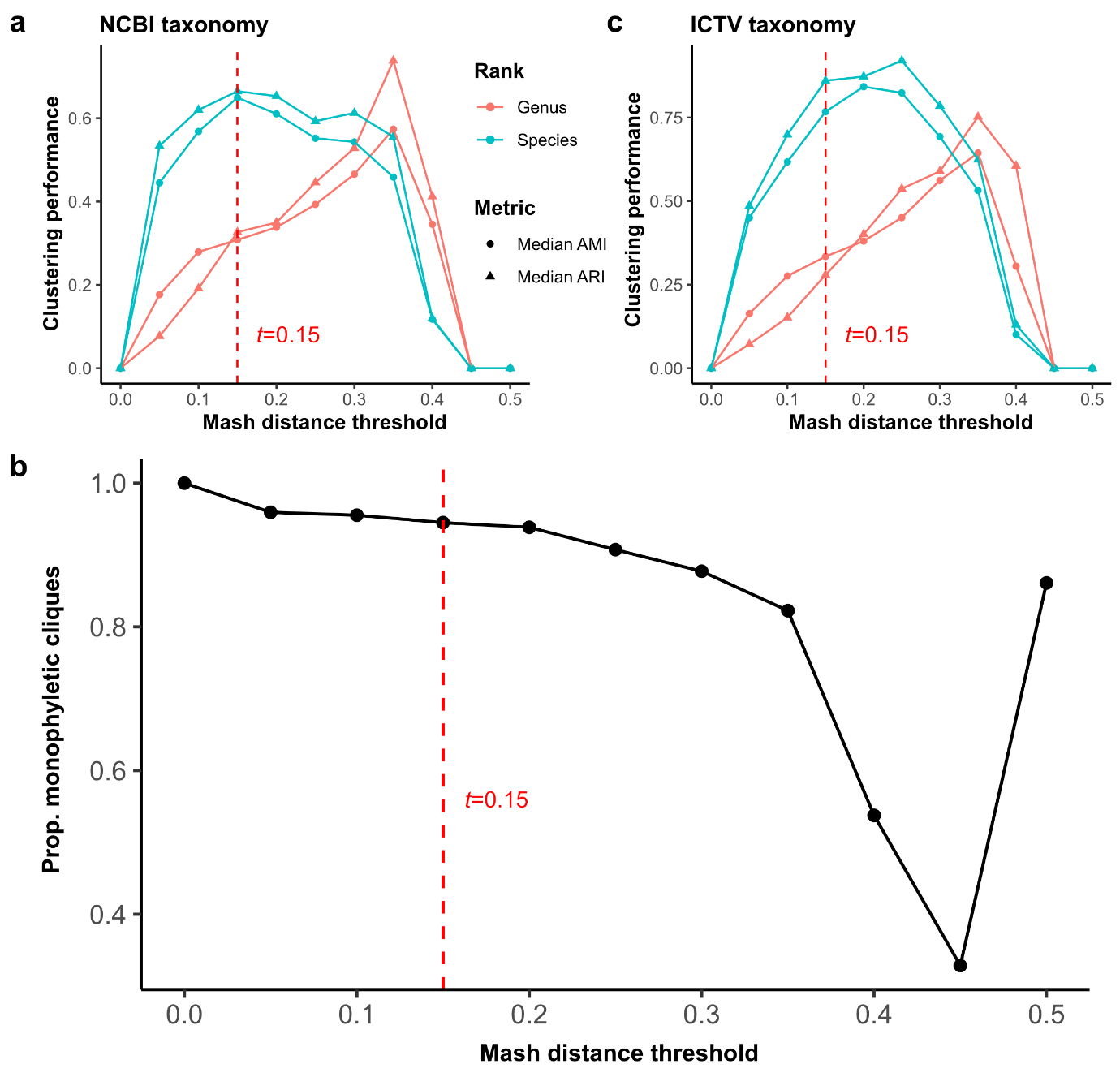

**Extended Data Figure 9**. **Selection of Mash distance threshold.** We constructed undirected, weighted graphs for each viral family with nodes and edges representing genomes and Mash distances, respectively. We then removed edges associated with values larger than the Mash distance threshold, *t*, before applying the community-detection algorithm, Infomap, to identify viral cliques. We then assessed the concordance of viral cliques identified against the (a) NCBI taxonomy or (c) ICTV taxonomy, using the median adjusted mutual information (AMI) or adjusted Rand index (ARI) across viral families. We also assessed (b) the overall proportion of monophyletic viral cliques across all viral families. The final distance threshold selected was *t*=0.15.

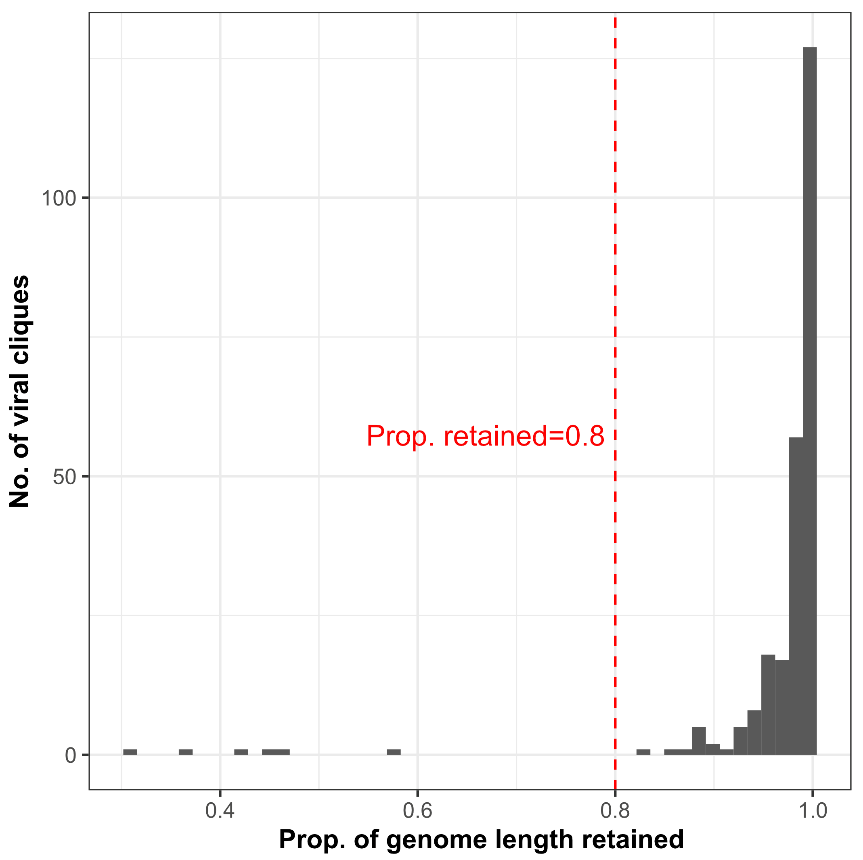

**Extended Data Figure 10. Assessing the quality of clique-level genome alignments.** For each viral clique, genomes were aligned and positions in these alignments were masked if more than 10% of the sequences corresponded to gaps or ambiguous nucleotides. The proportion of genome length retained was calculated as the number of unmasked sites divided by the median genome length of sequences in each clique-level alignment.

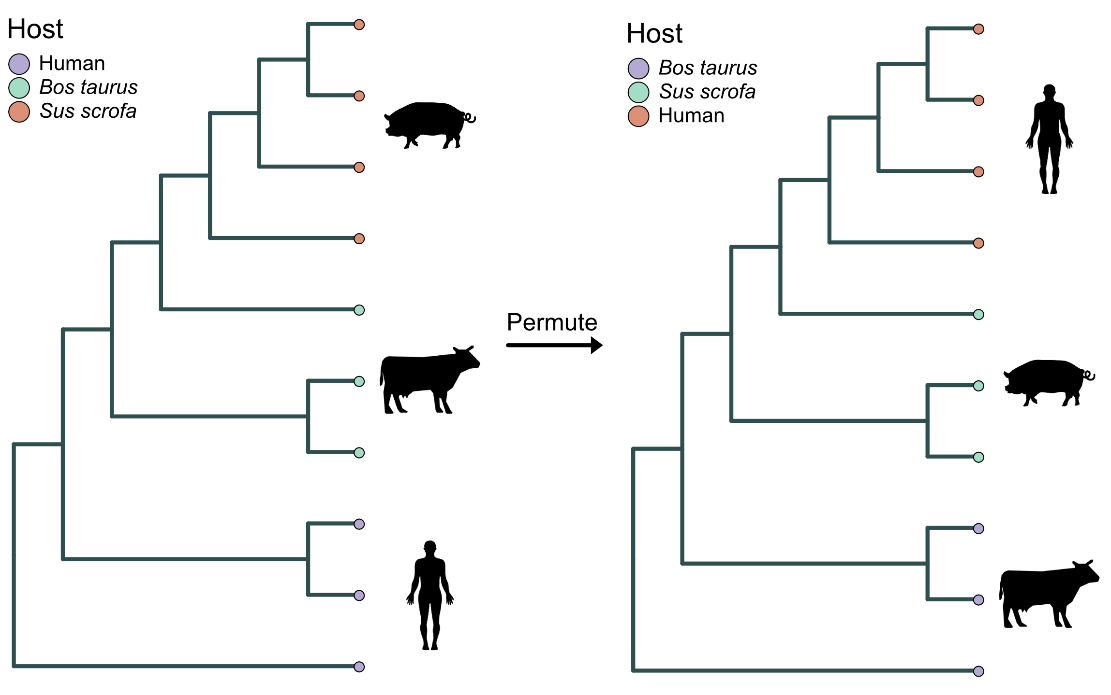

**Extended Data Figure 11**. Illustration of our permutation test approach applied to a hypothetical tree of a viral clique associated to three distinct hosts. Host states are permuted while retaining the number of host jumps inferred.

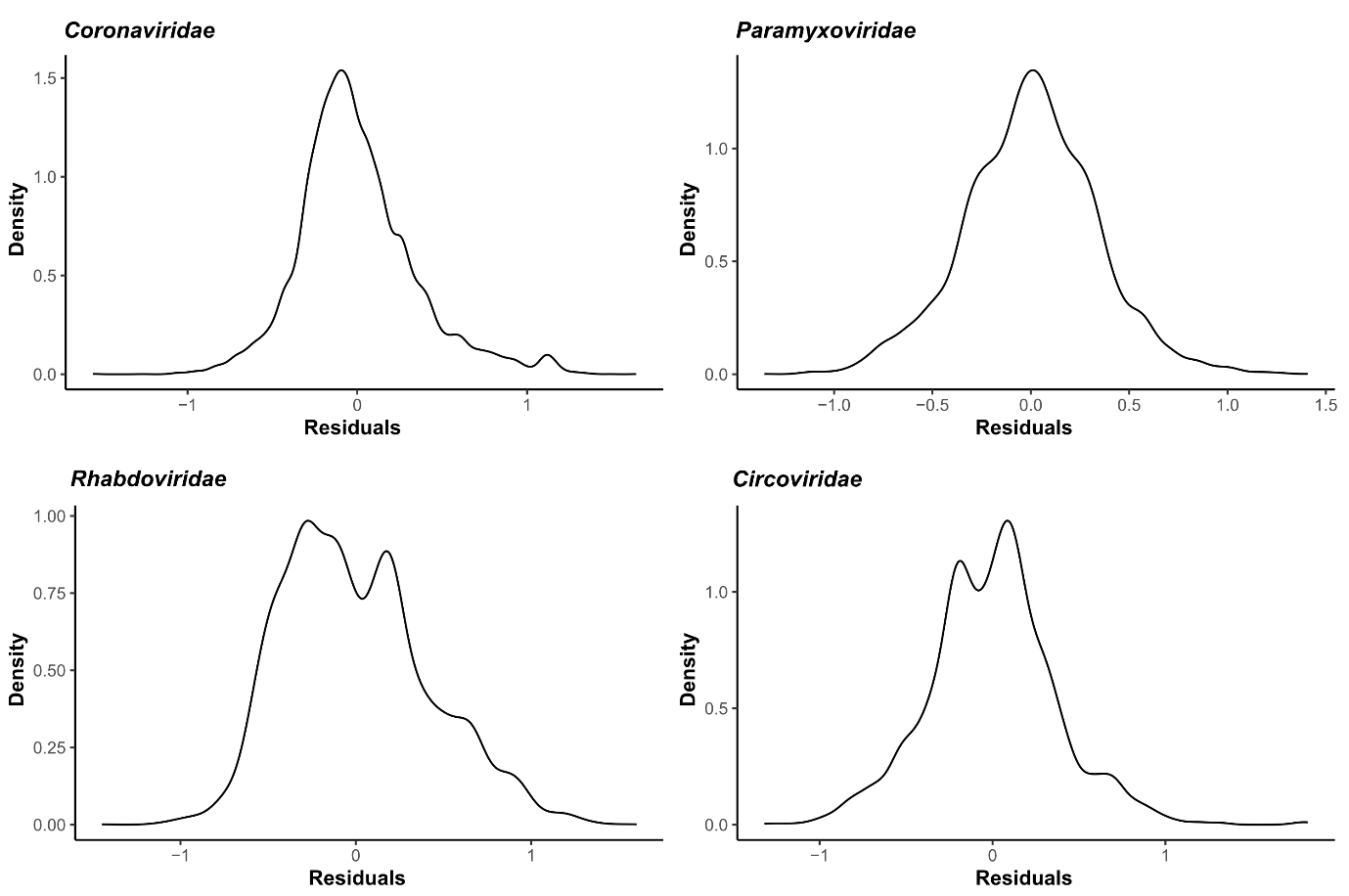

**Extended Data Figure 12. Linear regression model residuals.** Kernel density plots of residuals for the linear regression models implemented in Fig. 5.

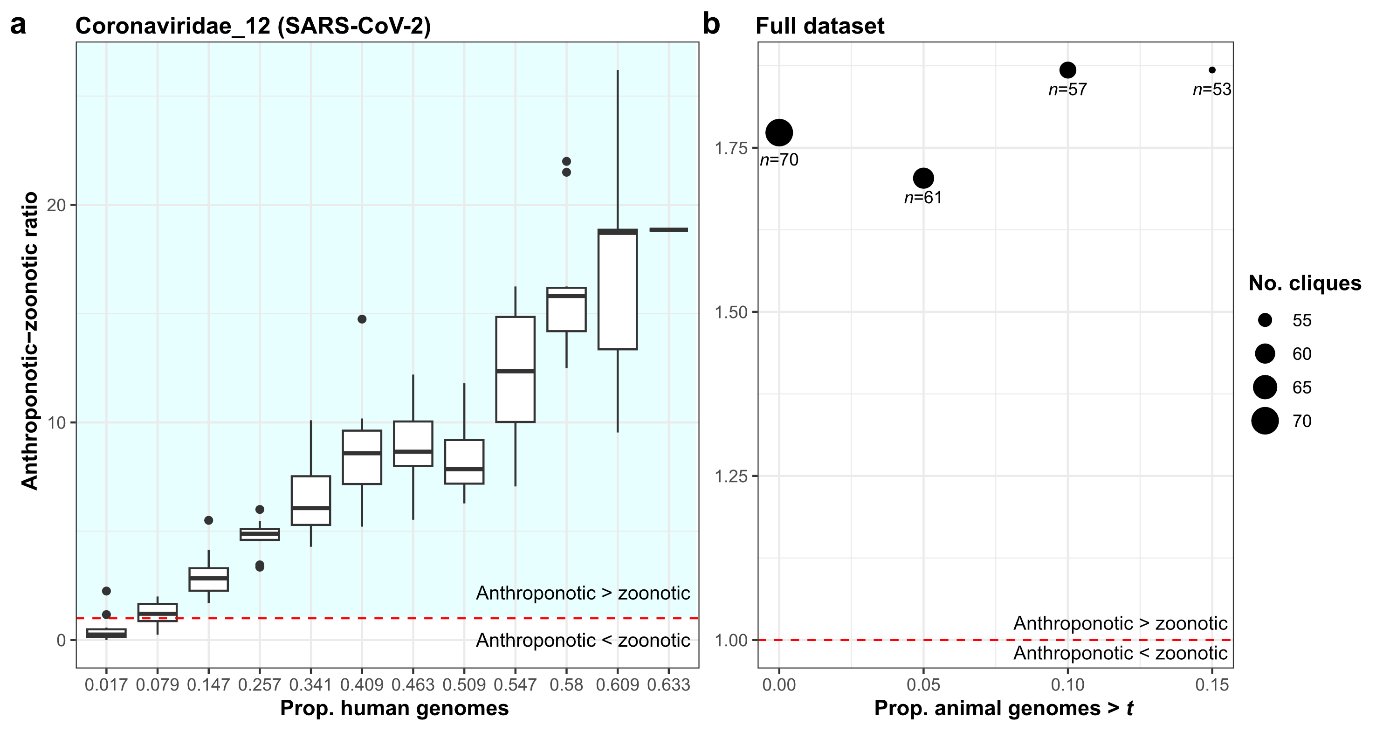

**Extended Data Figure 13**. (a) Effects of subsampling on the ratio of anthroponotic to zoonotic jumps inferred for Coronaviridae_12, the viral clique comprising SARS-CoV-2. Human-associated genomes were randomly subsampled to various degrees prior to ancestral reconstruction. A total of 10 iterations of the analysis per human-genome proportion were performed. Boxplot elements are defined as follows: centre line, median; box limits, upper and lower quartiles; whiskers, 1.5x interquartile range. (b) Ratio of anthroponotic to zoonotic jumps when retaining only viral cliques with greater than a certain proportion of animal-associated genomes, *t*, in the full dataset.

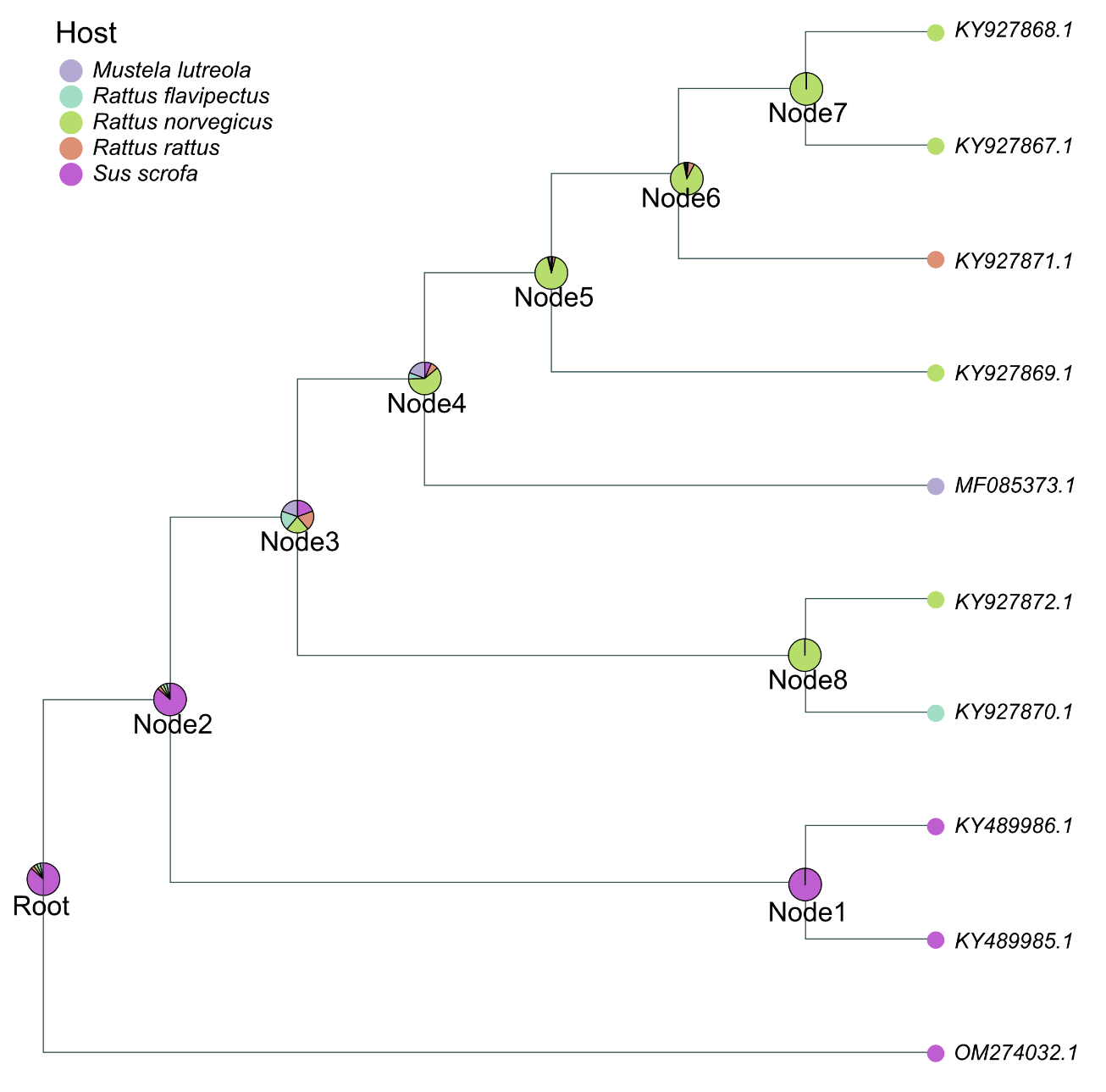

**Extended Data Figure 14**. Maximum likelihood tree for the viral clique Parvoviridae_58, rooted at OM274032.1. Piecharts show the relative likelihoods of all host states represented in this tree, as determined by maximum-likelihood ancestral state reconstruction.
